# Transfer learning of multicellular organization via single-cell and spatial transcriptomics

**DOI:** 10.1101/2024.02.28.582493

**Authors:** Yecheng Tan, Ai Wang, Zezhou Wang, Wei Lin, Yan Yan, Qing Nie, Jifan Shi

## Abstract

Spatial tissues exhibit complex gene expression and multicellular patterns that are difficult to dissect. Single-cell RNA sequencing (scRNA-seq) provides full coverages of genes, but lacking spatial information, whereas spatial transcriptomics (ST) measures spatial locations of individual or group of cells, with more restrictions on gene information. To integrate scRNA-seq and ST data, we introduce a transfer learning method to decipher spatial organization of cells named iSORT. iSORT trains a neural network that maps gene expressions to spatial locations using scRNA-seq data along with ST slices as references. iSORT can find spatial patterns at single-cell scale, identify key genes that drive the patterning, and infer pseudo-growth trajectories using a concept of SpaRNA velocity. Benchmarking on simulation data and comparing with multiple existing tools show iSORT’s robustness and accuracy in reconstructing spatial organization. Using our own new human artery datasets, iSORT shows its capability of dissecting atherosclerosis. Applications to a range of biological systems, such as mouse embryo, mouse brain, *Drosophila* embryo, and human developmental heart, demonstrate that iSORT can utilize both scRNA-seq and ST datasets to uncover multilayer spatial information of single cells.

## INTRODUCTION

Single-cell RNA sequencing (scRNA-seq)^1^ provides high-resolution and comprehensive transcriptomic profiles for all genes, allowing systematic analysis of cellular heterogeneity^2^, cell differentiation^3–5^, and disease mechanisms^6^. Computational analysis tasks of scRNA-seq data includes clustering^7^, cell types annotation^8^, differentially expressed gene identification^9^, and inferring pseudo-time^10^. Because the measured tissues are dissociated during the sequencing process, the information on the spatial locations of individual cells are lost in the scRNA-seq data.

Spatial transcriptomics (ST)^11^ can simultaneously capture information of both gene expressions and cell locations, providing a more desirable approach to study multicellular spatial organization. The two main types of ST technologies include the image-based and the sequencing-based. Image-based methods, such as MERFISH^12^, seqFISH^13^, and STARmap^14^, can detect only hundreds to thousands of genes at cellular or sub-cellular resolution. Sequencing-based methods, such as 10X Visium^15^ and Slide-seq^16, 17^, can provide whole transcriptomic sequencing, but only have a resolution at spot of group of cells instead of individual cells. Stereo-seq^18^ can capture thousands of genes in nanoscale resolution.

To utilize the strength of each data types, several computational tools were introduced to integrate scRNA-seq and ST data. For spot deconvolution, SPOTlight^19^ used the non-negative matrix factorization; Cell2location^20^ built a hierarchical Bayesian framework; Tangram^21^ and STEM^22^ employed deep neural networks on discrete spots. For estimating the single-cell location, novoSpaRc^23^ and spaOTsc^24^ used the optimal transport method to predict a spatial probability distribution for each individual cell; CSOmap^25^ estimated cell-cell affinity through ligand-receptor interactions; scSpace^26^ used a multilayer perceptron to extract features through transfer component analysis; CeLEry^27^ employed deep learning with enhancing ST by data augmentation; and CellTrek^28^ used a random forest approach with extensively interpolated ST data. Comparisons of those methods were carried out recently^29, 30^.

One major challenge for studying the multicellular organization using both scRNA-seq and ST data is to identify key genes that drive the spatial patterning of cells. Spatially Variable Genes (SVGs)^31^, which are genes with high spatial variability in expressions, may mark the spatial domains in gene expression pattern, however, they are not necessarily the genes that are responsible for the formation of the spatial patterns. Here we introduce a quantity named spatial-organizing genes (SOGs) based on the concept of dynamical causality^32, 33^. In other words, SOGs are characterized as the genes whose change critically affect the spatial organization of tissues.

The other major challenge is to demonstrate the differentiation trajectory of tissues in the physical space. RNA velocity proposed in 2018 used information of spliced and unspliced mRNA to derive a vector field in gene expression space presenting the direction of differentiation^34, 35^. Further generations include scTour^36^, which infer RNA velocity only by spliced mRNA expressions. However, RNA velocity does not consider the cellular organization and ignores the practical growth of cells in the physical space. Here we propose a quantity named SpaRNA velocity which projects the RNA velocity onto the ST slice, indicating pseudo-growth trajectories that model how cells transition to their progenies in space.

In order to estimate these two quantities, we utilize the density ratio technique in transfer learning by integrating large amount of scRNA-seq data with one or a few ST slices as references. In this **i**ntegrative method for **S**patial **O**rganization of cells using density **R**atio **T**ransfer (iSORT), a function using a neural network is constructed to connect the spatial organization of single cells and the gene expression. With this function, we can estimate the sensitivity of the spatial pattern with respect to individual genes as a measure to quantify SOG. Meanwhile, SpaRNA velocity is obtained by transferring RNA velocity to the physical space. To validate the effectiveness of iSORT, we collected benchmark datasets from human dorsolateral prefrontal cortex (DLPFC), mouse embryo, and mouse brain to test its accuracy and robustness in spatial reconstruction. *Drosophila* embryo dataset was used to show iSORT’s ability to reveal specific patterns. We conducted sequencing experiments to collect scRNA-seq and ST data from human arteries, in order to explore the potential influence of SOGs with atherosclerosis. SpaRNA velocity was visualized using DLPFC dataset and a human developmental heart dataset to illustrate pseudo-growth trajectories.

## RESULTS

### An overview of iSORT

iSORT is a transfer learning-based framework which constructs a neural network mapping gene expression to spatial coordinates and further analyzes the spatial organization (Fig. 1). One or several low-resolution ST slices are used as references in the training process. iSORT integrates scRNA-seq and ST data, which can be sampled from heterologous sources with different cell-type distributions. Estimation of the density ratio is the core technique. There are four major steps in the framework of iSORT, whose details are described in Methods. Specifically,

**Figure 1.**
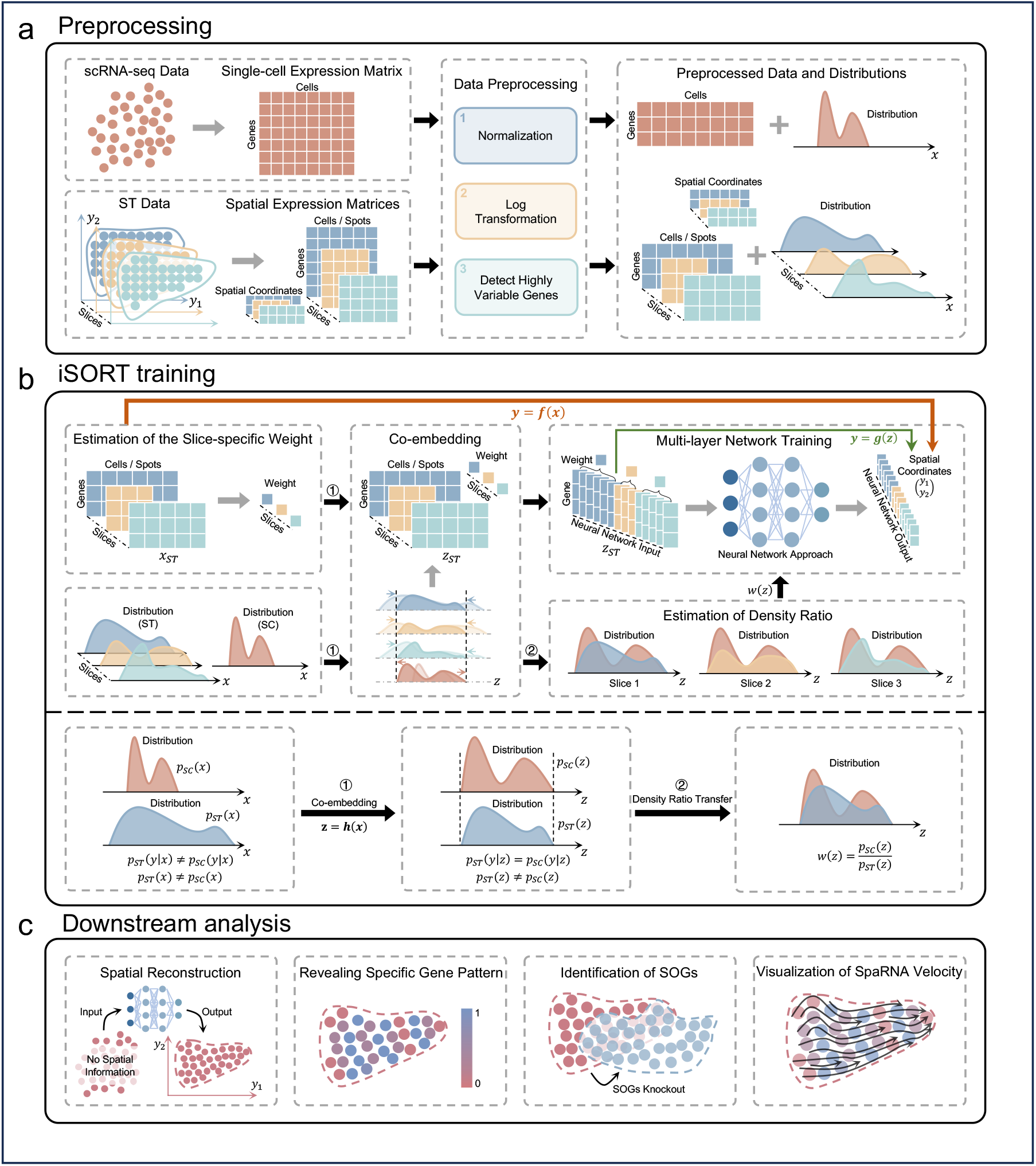
Overview of iSORT. (a) Preprocessing. Raw data from scRNA-seq dataset and ST slices are preprocessed and taken as iSORT’s inputs. Different samples could have diverse distributions of gene expressions. (b) Training. iSORT trains a mapping ***f*** = ***g*** ∘ ***h*** from gene expression ***x*** to spatial location ***y***, where ***z*** = ***h***(***x***) co-embeds data into the feature space in unified scale and ***y*** = ***g***(***z***) constructs a neural network combining slice-specific weights and density ratios. (c) Downstream analysis. By the mapping ***f***, iSORT can reconstruct spatial organization of tissues at single-cell resolution, reveal spatial expressive patterns of genes, identify SOGs, and visualize SpaRNA velocity in the physical space.

#### Step 1: Preprocessing (Fig. 1(a))

The inputs for iSORT, matched scRNA-seq data and ST slices, are preprocessed normalization, log transformation, and gene selection sequentially.

#### Step 2: iSORT training (Fig. 1(b))

Denote ***x*** as the gene expression after the preprocessing. Distributions of ***x*** in scRNA-seq and ST data are usually totally different. iSORT first employs a reference-based co-embedding approach^37^ to obtain a feature vector ***z***, which is marked as ***z*** = ***h***(***x***). Cells with similar features have close spatial coordinate ***y*** in the feature space, i.e. *p*_st_(***y*|*z***) = *p*_sc_(***y*|*z***), where *p*_st_ and *p*_sc_ are distributions of the scRNA-seq and ST data, respectively. Then, to transfer the spatial information from ST to the scRNA-seq data, we estimate the mapping ***y*** = ***g***(***z***) by minimizing a loss function *L*(***z, y***; ***g***), where the density ratio *w*(***z***) = *p*_sc_(***z***)/*p*_st_(***z***) measures the differences between scRNA-seq and ST data and is used for integrating the two types of data during training. In summary, iSORT constructs a mapping ***f*** = ***g*** ∘ ***h*** by neural networks, which assigns coordinates ***y*** = ***f*** (***x***) to each cell with gene expression ***x***.

#### Step 3: Downstream analysis (Fig. 1(c))

iSORT reconstructs the spatial organization of cells by mapping the gene expression ***x*** of each cell in scRNA-seq data to spatial coordinates ***y*** using the function ***f***. Results are naturally in single-cell resolution, and massive low-cost scRNA-seq data can be reorganized by only one or a few ST references. For the downstream analysis: (1) We reveal spatial patterns by analyzing the distribution and clustering of specific gene expressions within the spatial coordinates ***y***. (2) We identify SOGs by computing the index 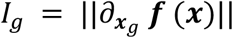, which quantifies the influence of each gene on the spatial structuring of the cell populations. (3) We define a quantity SpaRNA velocity as ***v***_st_ = ***f***(***x* + *v***_RNA_) **− *f***(***x***), where ***v***_RNA_ is the classical RNA velocity^36^ in the gene expression space. By integrating RNA velocity into our spatial model, one can analyze cellular dynamics on a ST slice, illustrating pseudo-growth trajectories of cells and elucidating how cells might migrate and evolve in physical space.

### Benchmarking iSORT in reconstructing the spatial organization of single cells

We first evaluated the performance of iSORT on reconstructing the spatial organization of three benchmark datasets, including the human dorsolateral prefrontal cortex (DLPFC) data, the spatially resolved mouse embryo data, and the mouse brain data. Details of the datasets are described in Supplementary Note 4.

### Reconstructing human DLPFC dataset

The DLPFC data comprises three post-mortem brain samples, with each sample containing four ST slices^38^. The ST data were obtained by the spot-size resolution 10X Visium technology. We took the ST slice ID151674 as the scRNA-seq input by removing its spatial coordinates. The ST slice ID151675 was used as the reference. The output of iSORT were compared with the ground truth (ST slice ID151674) as well as five other existing methods scSpace^26^, Tangram^21^, novoSpaRc^23^, CeLEry^27^, and CellTrek^28^ (Fig. 2(a)). Detailed information on the other five methods is shown in Supplementary Note 2. iSORT, Tangram, novoSpaRc, and CeLEry could reconstruct the shape of ST slice ID151674 from gene expressions, while iSORT, scSpace, novoSpaRc, and CeLEry could distinguish different layers with clear boundaries. Four different indicators were used to evaluate the performance of the six methods (Fig. 2(b)), i.e. the intra-layer similarity *S_C_*, the normalized density distribution *ρ_C_*, the aggregative volume index *A_C_*, and the aggregative perimeter index *P_C_* (Definitions and details are provided in Supplementary Note 5). Values of the indicators measure the performance of algorithms in reconstructing layer structures. iSORT got an average *S_C_* as 0.92, *ρ_C_* as 0.89, *A_C_* as 1, and *P_C_* as 1, which were larger than the values obtained from the other five methods (Fig. 2(b) and Supplementary Fig. 1).

**Figure 2.**
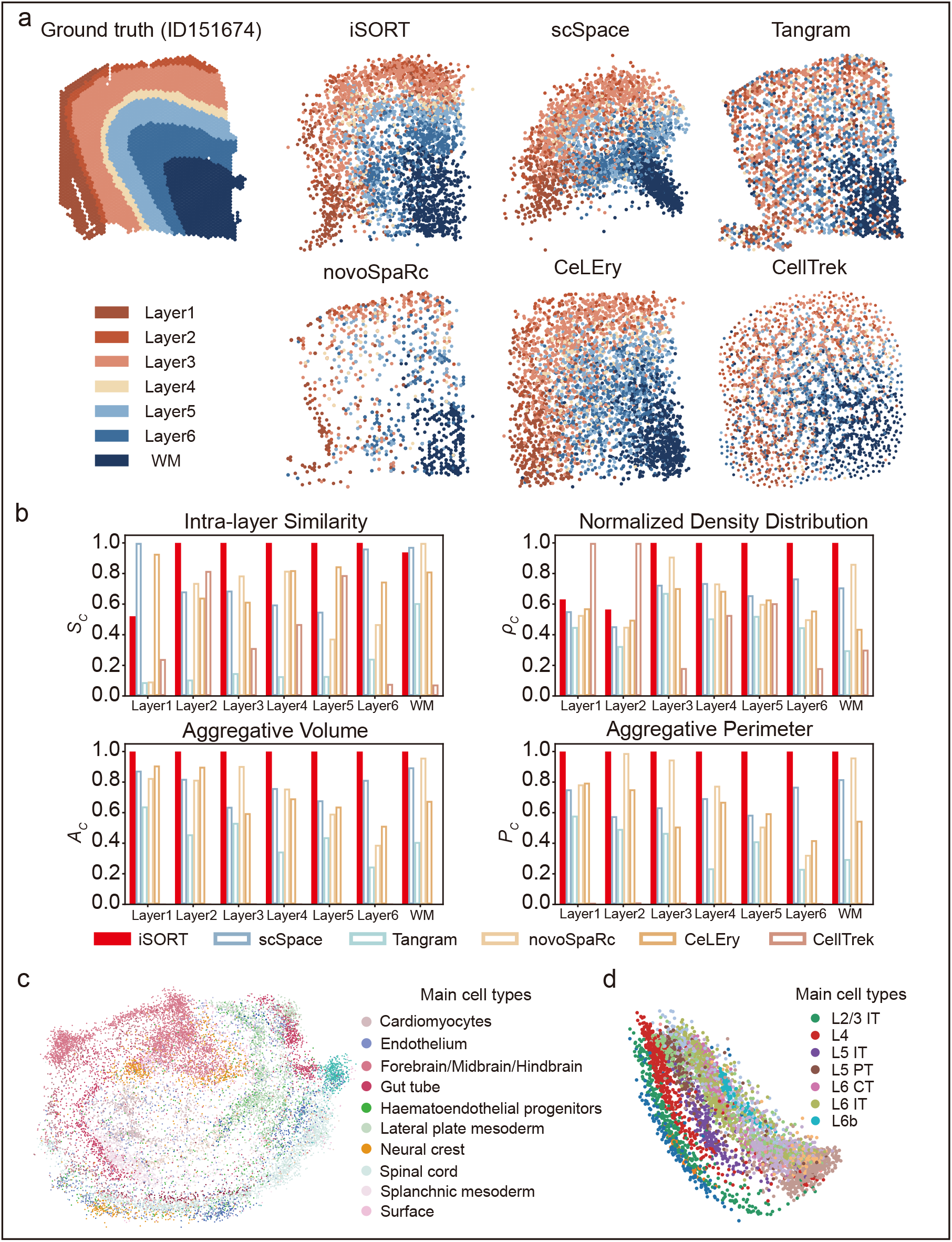
Spatial organization reconstruction by iSORT on the DLPFC, spatially resolved mouse embryo, and mouse brain datasets. (a) Ground truth and spatial reconstruction results for ID151674 in DLPFC dataset by six algorithms: iSORT, scSpace, Tangram, novoSpaRc, CeLEry, and CellTrek. Different colors represent different cortical regions. (b) Bar charts to compare the performance of six algorithms based on four indicators: intra-layer similarity (*S_C_*), normalized density distribution (*ρ_C_*), aggregative volume index (*A_C_*) and aggregative perimeter index (*P_C_*) of each layer. The overall average of indicators of iSORT (the red filled bars) is 0.89, larger than the other five algorithms. (c) Spatial reconstruction of the spatially resolved mouse embryo dataset with iSORT. Different colors represent different cell types. iSORT can reorganize single cells based on their gene expressions in a continuous space. (d) Spatial reconstruction of the mouse brain dataset with iSORT. iSORT can recover the stratified structures in the mouse brain.

### Reconstructing spatially resolved mouse embryo dataset

The mouse embryo data were sequenced by the seqFISH technology, involving sagittal sections at the 8-12 somite stage (E8.5-E8.75)^39^. The scRNA-seq input was taken as the gene expression from each spot by removing its spatial coordinates. As the spot of seqFISH reaches the single-cell resolution, in order to test the performance of iSORT on a low-resolution ST reference, we simulated a coarse-grained ST reference (details in Methods). The ground truth of the mouse embryo data is characterized by a rich diversity of cell types, which are displayed in a non-clustered pattern (Supplementary Fig. 2). The spatial reconstructions of scRNA-seq data by scSpace and CeLEry mixed different cell types together and were generally unable to capture the inner structure of the mouse embryo (Supplementary Fig.∼2). novoSpaRc and Tangram were constrained by the sparsity of the simulated spots and cells were only assigned to the spots (Supplementary Fig. 2). iSORT, however, reconstructed the spatial organization in a continuous space with single-cell resolution and retained the embryonic multicellular structure (Fig. 2(c) and Supplementary Fig. 2).

### Reconstructing mouse brain dataset

For the mouse brain datasets, the scRNA-seq and ST data were obtained from different samples, by different technologies, and with different cell-type distributions^40^. The scRNA-seq was conducted by Smart-seq2 and ST was sequenced by 10X Visium. Using iSORT, we reconstructed the spatial organization of cells in scRNA-seq data (Fig. 2(d)). iSORT reconstructed the stratified architecture of the cerebral cortex, which reflected the sequential arrangement of seven laminar excitatory neuron subgroups: L2/3 intratelencephalic (IT), L4, L5 IT, L5 pyramidal tract neurons (PT), L6 IT, L6 corticothalamic (CT), and L6b. scSpace and CeLEry hardly presented the stratified structures in the continuous space, while novoSpaRc and Tangram were limited by the spot resolution (Supplementary Fig. 3)

**Figure 3.**
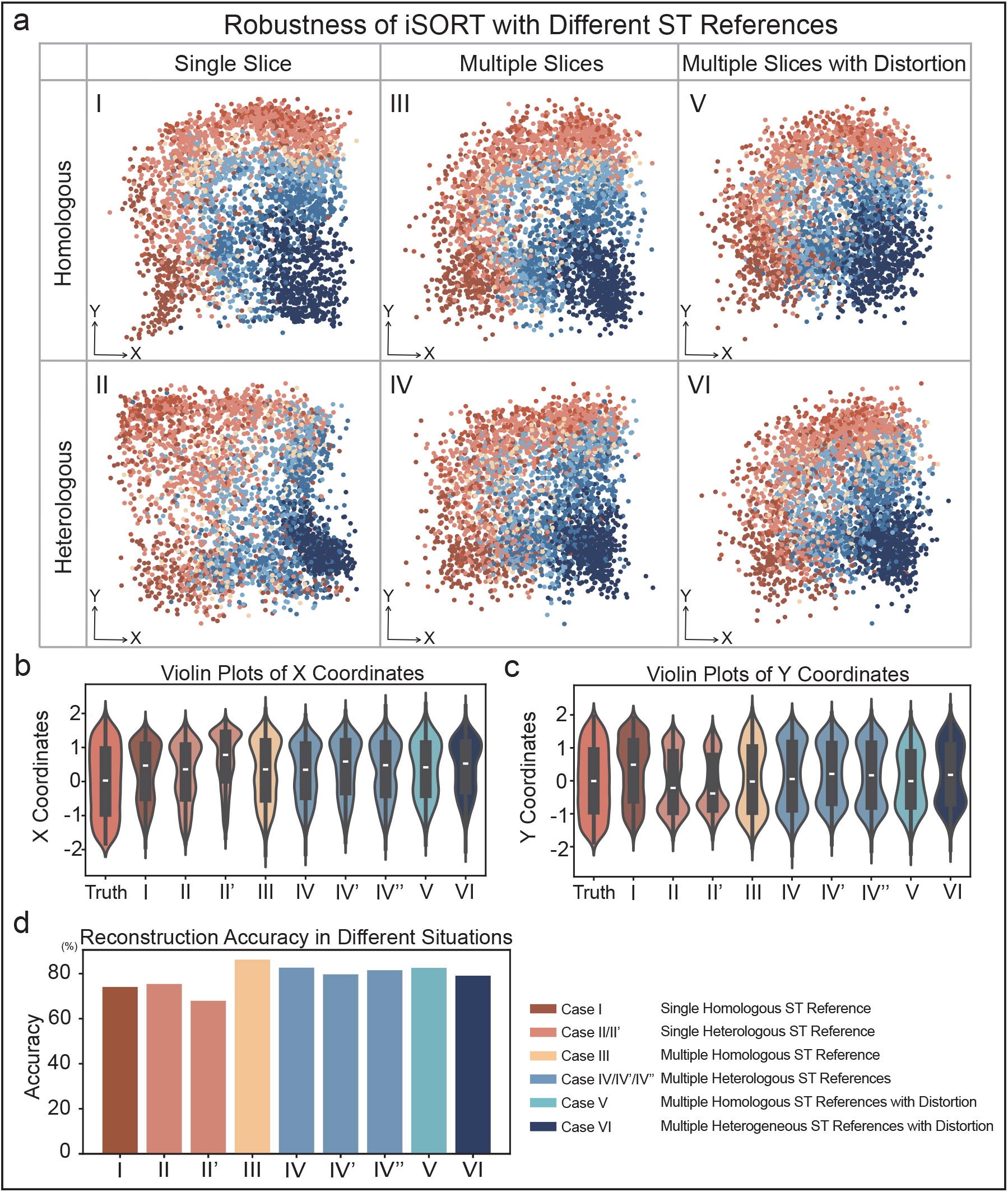
Robustness of iSORT in reconstructing human DLPFC slice ID151674 with different ST references. (a) iSORT’s results with different ST references. Case I: Reconstruction using one homologous slice (Sample ID151675). Case II: Reconstruction using one heterologous slice (Sample ID151671). Case III: Reconstruction using three homologous slices (Sample ID151673, ID151675, and ID151676). Case IV: Reconstruction using three heterologous slices (Sample ID 151675, ID151607, and ID151671). Case V: Reconstruction using three rotated homologous slices (Sample ID151673, ID151675, and ID151676). Case VI: Reconstruction using three rotated heterologous slices (Sample ID151675, ID151671, and ID151507). (b) Violin plots of X coordinates across different reconstruction scenarios, depicting the distributions of cells on the X-axis. Case II’: Reconstruction using one heterologous slice (Sample ID151570). (c) Violin plots of Y coordinates across different reconstruction scenarios, depicting the distributions of cells on the Y-axis. Case IV’: Reconstruction using three heterologous slices (Sample Id151675, ID151507, and ID151508). Case IV”: Reconstruction using three heterologous slices (Sample ID151675, ID151670, and ID151671) (d) The accuracy of iSORT across different cases.

### Robustness of iSORT in integrating spatial information from different slices

The ST slices can be sampled from heterologous data sources, with diverse cell-type distributions, and being sequenced under different structural distortions. To study how sensitive iSORT is when a variety of ST slices are used as references, we systematically tested the human DLPFC dataset. The same ’scRNA-seq’ data (ST ID151674 with the spatial information removed) was used to reconstruct its spatial structure. Original ST slices of DLPFC from different samples used in the sensitivity test showed various hierarchical distributions (Supplementary Fig. 4). Six cases using different ST references were studied systematically (Fig. 3(a)).

**Figure 4.**
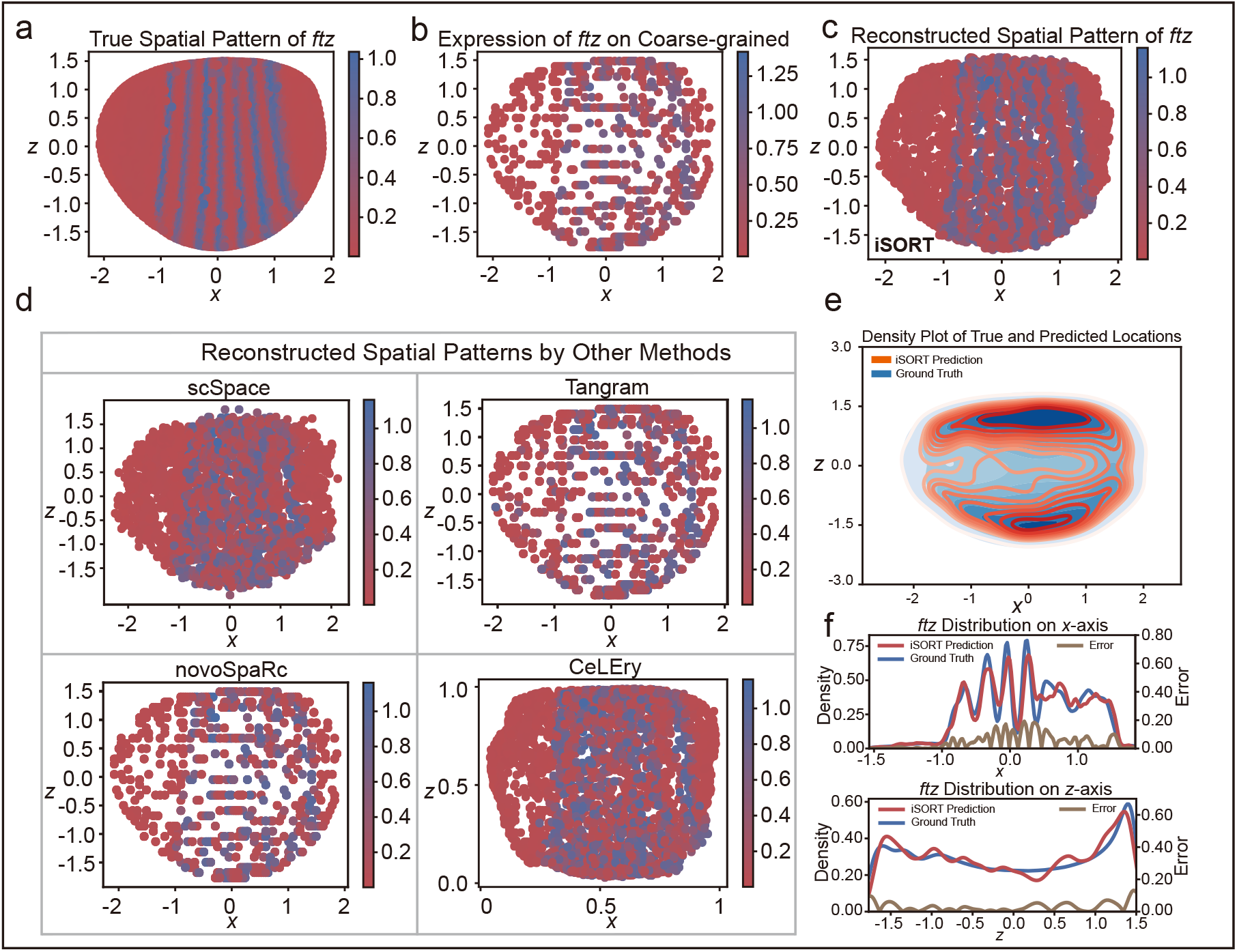
Revealing the spatial pattern of the *ftz* gene from Drosophila embryo data using iSORT. (a) Visualization of *ftz*’s spatial pattern in the original ST slice. (b) Visualization of *ftz*’s expression in the simulated coarse-grained ST reference, demonstrating the disruption of gene patterns in the low-resolution ST spots. (c) Visualization of *ftz*’s spatial pattern reconstructed from the scRNA-seq data by iSORT. (d) Reconstructed spatial patterns by scSpace, Tangram, novoSpaRc and CeLEry. (e) Density plot contrasting the true (blue) and predicted (red) spatial locations of cells. (f) Marginal densities for the true (blue) and predicted (red) spatial distributions of the *ftz* gene. The errors between the true and predicted densities are shown by the yellow line. The seven stripes along the x-axis are recovered by iSORT.

#### Case I: Single homologous ST reference

The single ST reference ID151675 was from the same DLPFC region as the scRNA-seq (ID151674). The reconstruction of iSORT (Fig. 3(a-I)) is evaluated by four clustering indicators (Fig. 2(b)). We also measured the accuracy of the reconstruction (Supplementary Note 5), which reaches 100% for an exact spatial reconstruction but 0% for a random guess. iSORT obtains an accuracy of 74.1% for the case with a single homologous ST reference (Fig. 2(d) and Supplementary Table 1).

#### Case II: Single heterologous ST reference

The single ST reference ID151671 was from a heterologous tissue. iSORT obtained a spatial reconstruction with an accuracy of 75.4% (Figs. 3(a-II) and (d)). Another heterologous reconstruction using ID151507 (Case II’) as the reference got an accuracy 68.0% (Supplementary Fig. 5). In Case II and II’, structural layers of the target sample were recovered as well as Case I.

**Figure 5.**
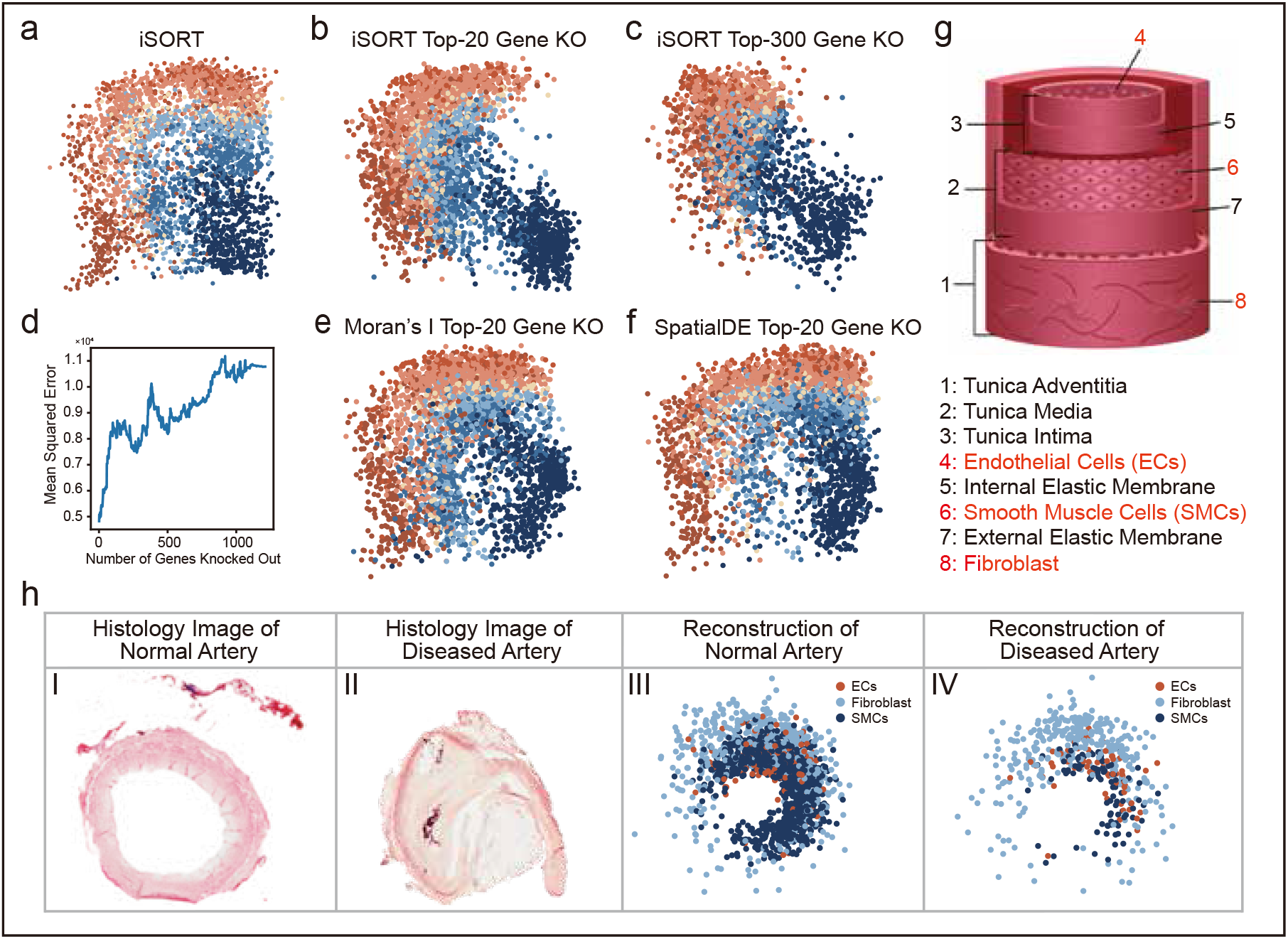
In-silico gene knockout experiments on DLPFC dataset and analysis of the human artery dataset. (a) Reconstruction of DLPFC by iSORT without gene knockout. (b) Reconstruction of DLPFC with top-20 SOGs knocked out (KO). (c) Reconstruction of DLPFC with top-300 SOGs knocked out. (d) Curve of the mean squared error (MSE) in reconstruction with increasing SOGs knocked out. (e) Reconstruction of DLPFC with the top-20 Moran’s *I* SVGs knocked out. (f) Reconstruction of DLPFC with the top-20 SpatialDE SVGs knocked out. (g) Schematic diagram of artery structure illustrating layered composition: the innermost layer is lined with endothelial cells (EC), followed by smooth muscle cells (SMC), and the outermost layer composed of fibroblasts. (h) Hematoxylin and Eosin staining and the reconstruction results of the normal artery and the diseased artery with atherosclerosis (AS): Panel I: Histology image of a normal artery. Panel II: Histology image of an artery with AS. Panel III: Reconstruction result for a normal artery, showcasing ECs, SMCs, and fibroblasts. Panel IV: Reconstruction result for a diseased artery with AS, showcasing ECs, SMCs, and fibroblasts.

#### Case III: Multiple homologous ST references

In this case, we used three homologous slices (ID151673, ID151675, ID151676) as the ST references. iSORT achieved a higher accuracy of 86.2% in reconstructing spatial organization than Case I, which is the highest value among all six cases (Figs. 3(a-III), (d), and Supplementary Table 1). The results implied that using multiple ST references could improve the accuracy of iSORT.

#### Case IV: Multiple heterologous ST references

In practice, scRNA-seq data and ST slices usually originate from different samples with different shapes and cell-type distributions. In this case, we used slices ID151507, ID151675, and ID151671 as the ST references. Although the diversity and complexity between the scRNA-seq and ST data were high, iSORT still reconstructed the hierarchical structure with an accuracy of 82.6% (Fig. 3(a-IV)). For other combinations of heterologous references, we presented Case IV’ using slices ID151675, ID151507, and ID151508 with an accuracy of 79.7%, and Case IV” using slices ID151675, ID151670, and ID151671 with an accuracy 81.5% (Supplementary Fig. 5).

#### Case V: Multiple homologous ST references with distortion

To further test the robustness of iSORT, we added rotations to the ST references, which is one of the most common batch effects of experimental data. Homologous ST slices ID151673, ID151675, and ID151676 as in Case III were rotated by 45, 0, and **−**45 degree, respectively (Supplementary Fig. 6). The accuracy value 82.6% (Fig. 3(a-V)) showed that iSORT could produce spatial reconstruction even when there were noticeable distortions in the input slices.

**Figure 6.**
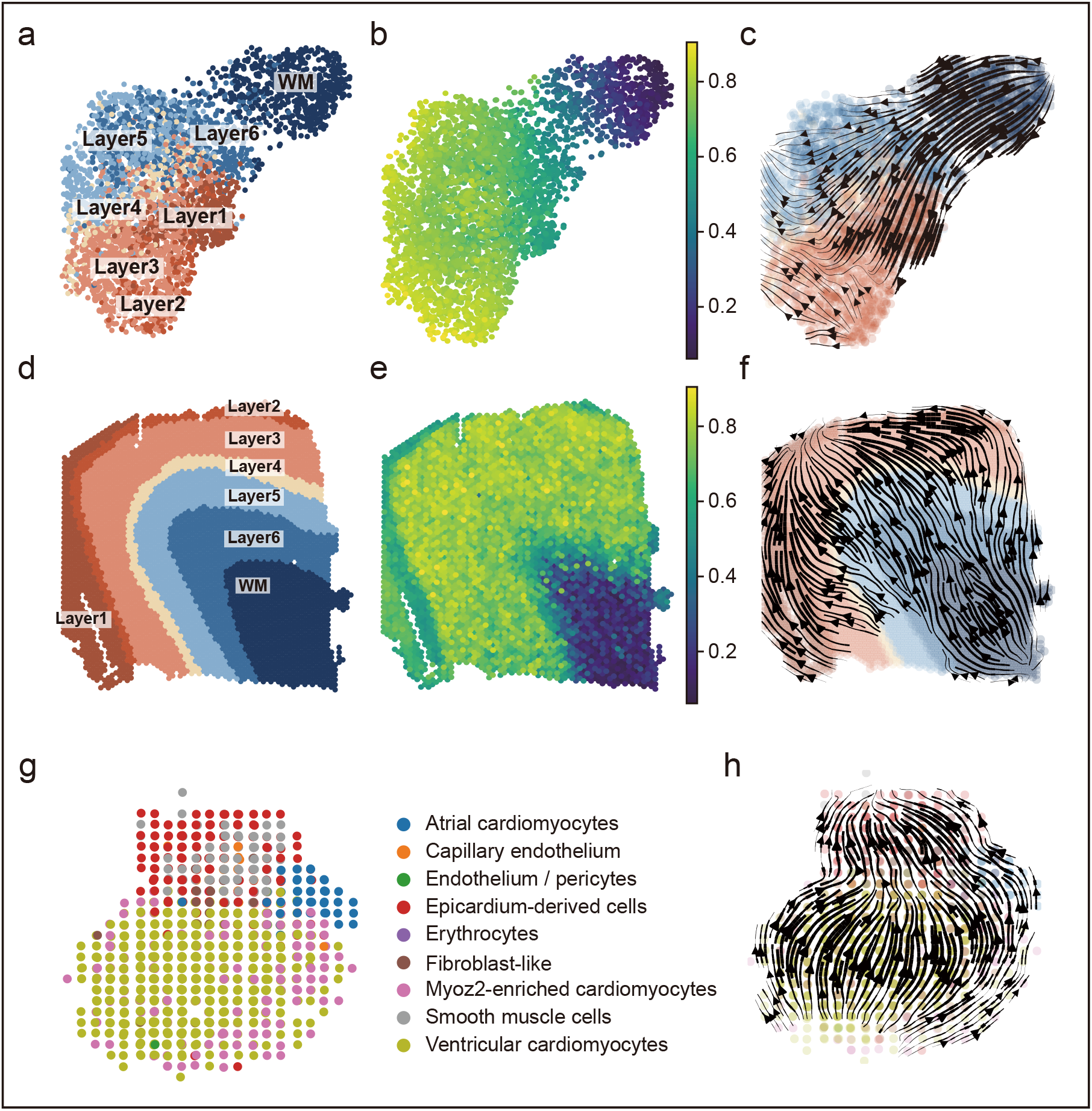
Visualization of SpaRNA velocity on DLPFC and human developing heart dataset. (a) Sample ID151674 shown in UMAP-reduction gene expression space annotated by different layers. (b) Pseudo-time of sample ID151674 inferred by scTour shown in UMAP-reduction gene expression space. (c) RNA velocity of sample ID151674 inferred by scTour shown in UMAP-reduction gene expression space. (d) Sample ID151674 shown in physical space annotated by different layers. (e) Pseudo-time of sample ID151674 shown in physical space. (f) SpaRNA velocity of sample ID151674 inferred by iSORT shown in physical space. (g) Spatial transcriptomics of a human developing heart at 9 post-conception week. (h) SpaRNA velocity on the human developing heart visualized by iSORT.

#### Case VI: Multiple heterogeneous ST references with distortion

The heterogeneous ST slices ID151507, ID151675, and ID151671 as in Case IV were rotated by 30, 0, and **−**40 degree, respectively (Supplementary Fig. 6). iSORT’s reconstruction captures the spatial organization with an accuracy of 79.1% (Fig. 3(a-VI)).

We also studied the spatial distributions of the true cell positions and reconstructions in different cases by violin plots (Figs. 3(b-c)). An index of accuracy (Supplementary Note 5) was computed for comparing different cases, where Case III held the largest similarity (highest accuracy) with the ground truth (Fig. 3(d)). iSORT obtained an accuracy over 74.0% in most cases (excepts Case II’ with 68.0%), which quantitatively demonstrated that iSORT was able to robustly integrate the DLPFC spatial information from multiple ST slices under various conditions (Fig. 3(d) and Supplementary Table 1).

### Uncovering spatial strip patterns of gene expression in *Drosophila* embryo

To study how iSORT uncovers the spatial organization of gene expression pattern, we applied iSORT to the *Drosophila* embryonic development dataset^41, 42^. The ST data was sequenced by FISH-seq. The scRNA-seq input was obtained by removing the spatial information, while a low-resolution ST reference was obtained by the coarse-grained simulation (Fig. 4(b) and more details in Methods). During the early development of embryo, several genes, such as *ftz*, exhibit unique spatial patterns and play decisive roles in the axial establishment and segmentation of the *Drosophila* body^43^. As an example, *ftz* in the original ST slice showed a unique seven-striped spatial pattern (Fig. 4(a)). iSORT’s reconstruction showed the seven stripes, each corresponding to a future body segment (Fig. 4(c)). The true location distributions of single cells with the predicted ones by iSORT were compared, and iSORT captured the cell density of the embryo (Figure 4(e)). The marginal densities and corresponding errors between the true *ftz* expression and the predicted ones by iSORT were calculated, with reconstructed marginal densities on the x-axis showing the seven-striped pattern of *ftz* (Fig. 4(f)). We also compared with the reconstructions by scSpace, Tangram, novoSpaRc, and CeLEry (Fig. 4(d)). Tangram and novoSpaRc failed to fully recover the seven-striped pattern, due to the discrete low-resolution constraints. scSpace and CeLEry could not clearly separate different stripes or determine the stripe number. Furthermore, we tested iSORT on 12 other genes, including *Antp, cad, Dfd, eve, hb, kni, Kr, numb, sna, tll, twi*, and *zen*. The coarse-grained references (Supplementary Fig. 8) could not reflect the true spatial patterns of these 12 genes (Supplementary Fig.7), while the reconstructions by iSORT can reveal the corresponding patterns (Supplementary Fig.9). These results supported that iSORT could reveal spatial patterns of gene expression with good consistency with the original spatial organization.

### SOG analysis and identification of atherosclerosis-related biomarkers in human artery experiments

SOGs offer an approach to identify key genes determining the cellular spatial organization within tissues. Since iSORT provides a mapping ***y*** = ***f*** (***x***) from the gene expression ***x*** to spatial coordinates ***y***, the SOG index 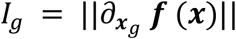 can measure the influence of gene *g* (see Methods). Genes with larger *I_g_* s have more impact on the spatial organization. The top genes with large *I_g_* values are defined as SOGs. To study SOGs, we analyzed two different datasets.

### SOGs in human DLPFC dataset and in-silico knockout validations

We chose ID151674 (removing spatial information) as the scRNA-seq input and ID151675 as the ST reference. With all 3635 genes, iSORT’s reconstruction distinguished the hierarchical structure of different cerebral cortexes (Figure 5(a)). After ranking the genes by the SOG index *I_g_*, we conducted the knockout validation. When the top 20 SOGs (Supplementary Table 2) were removed, the reconstructed spatial organization was significantly altered (Fig. 5(b)). When the top 300 SOGs were removed, the spatial structure was further affected (Fig. 5(c)). We calculated the mean square error (MSE) between the true cell locations and the reconstructed ones. MSE value was found to increase with the number of knockout SOGs (Fig. 5(d)). Moreover, we conducted the top-20-knockout experiments for SVGs selected by Moran’s *I* score^44^ and SpatialDE^31^ (Supplementary Tables 3 and 4). Moran’s *I* score assessed the spatial correlation and SpatialDE leveraged a Gaussian-process-based model on spatial expression (details in Supplementary Note 3). It was found that the spatial structures after knocking out SVGs were not significantly disrupted (Figs. 5(e-f)) compared to the SOGs (Fig. 5(b)). The results indicated that SOGs selected by iSORT instead of SVGs were the key genes to maintain tissue’s spatial organization.

### Human artery sequencing and analysis of atherosclerosis-related SOGs

Diseases like atherosclerosis (AS) may involve changes in spatial morphological features like subendothelial lipid deposition, narrowing of the vascular lumen, and thickening of the arterial wall^45^. Identification of disease-related SOGs can provide instructive information for the treatment^46^. To validate the potential of SOGs, we sequenced a new dataset from human diseased arteries with AS and normal arteries. We performed scRNA-seq on Illumina NovaSeq platform and obtained ST by 10X Genomics. We also conducted hematoxylin and eosin staining to show that diseased arteries appear to be blocked while the normal artery was in a circular shape (Figs. 5(h-I) and (h-II)). The detailed information of data collection and sequencing can be found in Methods section.

Because the artery is not a simple connected region but a circular shape (Fig. 5(g)), we applied a reversible polar transformation to the ST data in preprocessing before using iSORT to reconstruct the spatial organization (Supplementary Note 6). There were two independent scRNA-seq samples as inputs: one from an individual without AS, and the other from a patient with AS. Three ST slices were used as multiple heterologous references: two from individuals without AS, and one from a patient with AS. The scRNA-seq and ST slices were sampled from totally different sources. The iSORT reconstructions for the normal artery and the diseased artery with AS showed the correct sequence of layers from inside to outside: endothelial cells (ECs), smooth muscle cells (SMCs), and fibroblasts (Figs. 5(h-III) and (h-IV)). More detailed reconstructions for all cell types are exhibited in Supplementary Figs. 13 and 14. Compared with the normal artery (Fig. 5(h-III)), the spatial organization of the diseased artery with AS was more concentrated on one side of the vessel (Fig. 5(h-IV)). Accumulation of lipids and fibrosis within the vessel wall in patients with AS, makes more cells concentrate on one side and lead to hardening and narrowing of the artery^45^. Then, we ranked and selected top SOGs of normal and diseased arteries based on *I_g_* (Supplementary Fig. 10, Supplementary Tables 5 and 6). We found 16 important overlapping SOGs. Among them, *TNN* achieved the highest *I_g_* score, which plays a crucial role in facilitating integrin binding and is essential for cell adhesion, migration, and proliferation^47^. *TNN* is also important in neuronal generation and osteoblast differentiation, further underscoring its significance in vascular structure and function^48, 49^. Meanwhile, a number of genes found by *I_g_*, such as *RFLNA, SPN*, and *EZH2*, are proved to be associated with AS by literatures^50–52^. We further performed the GO enrichment analysis on the top-50 SOGs. The GO analysis revealed that 11 out of the top 20 results in GO Biological Processes (BP) were common to both the normal and AS samples (Supplementary Tables 7-8, and Supplementary Figs. 11-12). The shared BP terms, such as leukocyte proliferation and regulation of cell-cell adhesion, indicated commonality in core biological functions, while the unique BP terms appeared in the AS sample were highly related to the abnormal vascular mineralization during AS process. More detailed analysis of the AS related pathology can be found in Supplementary Note 8. These results indicated that the SOGs identified by iSORT contributed to maintaining vascular function and could serve as biomarkers for the study of AS.

### SpaRNA velocity and pseudo-growth trajectories on human DLPFC and developmental heart dataset

Next, we study the iSORT derived SpaRNA velocity and its effect on the tissue organization using two datasets.

### SpaRNA velocity on human DLPFC dataset

ST slice ID151674 was used in this experiment, with UMAP showing the distribution of cells (Fig. 6(a)). It was known that the growth starts from the white matter (WM) to the Layer 1, passing through Layer 6, Layer 5, Layer 4, Layer 3 and Layer 2^48, 53, 54^. The pseudo-time and RNA velocity inferred by scTour^36^ in the reduced two-dimension space could not capture the correct growth of Layer 1 (Figs. 6(b) and (c)). Pseudo-time inferred from the gene expression space exhibited discontinuities on the ST slice, such as between Layer 2 and Layer 3 (Figs. 6(d-e)). However, the SpaRNA velocity derived from iSORT reconstructed the correct growth trajectories from WM to the sequential layers. Moreover, iSORT also allows visualization of the detailed internal growth within a layer, particularly in the WM layer (Fig. 6(f)).

### SpaRNA velocity on human developmental heart dataset

Next, we presented the SpaRNA velocity derived by iSORT on the human developmental heart dataset^55^. This dataset contained ST slices from three developmental stages at 4.5-5, 6.5, and 9 post-conception weeks (PCWs). ST in the spot-size resolution with annotated cell types at 9 PCW showed the structure of a heart (Fig. 6(g)). The initial development involves ventricular myocardial cells, preceding cells located in the atrial region, including atrial cardiomyocytes, epicardium-derived cells, smooth muscle cells, and fibroblasts, which is in accordance with the SpaRNA velocity inferred by iSORT (Fig. 6(h)). Results for cells at 4.5-5 PCW and 6.5 PCW also showed similar pseudo-growth trajectories (Supplementary Fig. 15). The chronological progression of heart development in physical space, in line with established biological processes^56, 57^, can be quantitatively described by the pseudo-growth trajectories obtained from SpaRNA velocity.

## DISCUSSION

In this study, we introduce a comprehensive analysis tool for exploring and deciphering the spatial organization of cells. iSORT leverages the concept of density ratio transfer to integrate scRNA-seq data with one or more ST references and reconstruct the spatial structure of scRNA-seq data at single-cell resolution. By using the reconstruction mapping, iSORT can recover spatial patterns of gene expressions, identify SOGs that are critical in driving spatial organization, and introduce a new quantity SpaRNA velocity that captures pseudo-growth of tissue to model growth dynamics of tissue. iSORT is found to be robust under different situations, such as diverse sample sources and additional distortion of references. We also conducted sequencing experiments on human arteries with and without atherosclerosis to exhibit the practicality of iSORT in finding biomarkers and researching diseases.

Several areas of improvements can be addressed for iSORT.

First, during the training process by iSORT, ST references and scRNA-seq data are required to be the same or similar tissues, although their sampled objects, tissue shapes and cell-type distributions may be different. Allowing the training on more samples including different organs and species could significantly improve the power of the method. As large models in the single-cell domain are developing rapidly, models like scGPT^58^ and scFoundation^59^ demonstrate the feasibility and benefits of using large-scale pretrained models in understanding complex biological data. The iSORT framework, with its unique focus on integrating single-cell and ST data, could be ideally positioned to leverage these advancements. By incorporating elements from these large models, iSORT could enhance its capabilities in reconstructing spatial organization from diverse tissue types and conditions.

Second, cell-cell communication (CCC) inference provides a powerful way in analyzing spatial organization of tissue^60, 61^. With the spatial information inferred by iSORT on cells, one may use matched scRNA-seq data and ST data for CCC inference. In particular, CCC may be scrutinized in conjunction with SOGs to analyze patter-driven CCC.

Meanwhile, many diseases can be attributed to the abnormal spatial organization. Beyond selecting highly variable genes from the expressions, SOGs fully utilize the spatial information and provide a way to detect pathogenic genes. Besides the atherosclerosis described in this study, iSORT could offer a new perspective for revealing broader disease-related pathways.

The SpaRNA velocity derived by iSORT could reflect the organizing and differentiation process of tissues. It is based on the concept of RNA velocity to plot the pseudo-growth trajectories on ST. Other methods of RNA velocity, such as scVelo^62^, can also be appropriately modified for the SpaRNA velocity inference. Besides, the inference of pseudo-time is another important aspect in assessing cell developmental status^63, 64^. Inference of SpaRNA velocity directly from ST data could be helpful in the research of pseudo-time.

Recently, several methods have been proposed for joint 3D reconstruction of biological tissues using multiple 2D slices^65^. The framework of iSORT, in principle, can be generalized to incorporate 3D spatial reconstruction and the pseudo-growth dynamics.

In summary, iSORT provides a promising computational framework for analyzing and integrating scRNA-seq and ST data, offering perspectives in analyzing patterns and organization of spatial tissues.

## METHODS

### Data preprocessing

First, the raw gene expression for each scRNA-seq cell or ST spot is log-transformed and normalized by

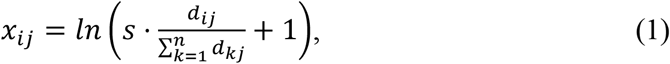

where *d*_*ij*_ is the raw expression of gene *i* in cell/spot *j, n* is the total number of genes, *s* is a scaling factor with default value 10000, and *x*_*ij*_ is the normalized gene expression. Then, *x*_*ij*_ is scaled by *z*-score

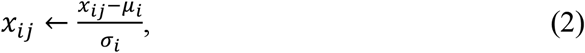

where *μ_i_* and *σ_i_* are the mean value and standard variation of gene *i* across cells/spots, respectively. Highly variable genes are selected for further studies. All these preprocessing procedures can be realized by Python codes or packages such as Seurat^37^ and Scanpy^66^.

### iSORT framework

In this section, we describe the framework of iSORT for the single ST reference case. Details for the case with multiple ST references can be found in Supplementary Note 1.

#### Spatial organization mapping

Denote 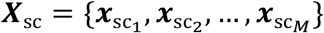 as the expressions for the scRNA-seq data, and 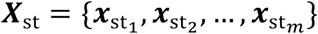 for the ST reference, where *M* and *m* are the sample sizes. The location of ST data is 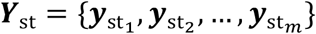. Suppose that there are *H* highly variable genes after preprocessing, then we have 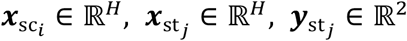, *i* = 1,2, …, *M, j* = 1,2, …, *m*. Let *p*_sc_(***x, y***) and *p*_st_(***x, y***) be the joint probability density functions (pdfs) of the scRNA-seq data and the ST data, respectively. Due to the variety in samples and discrepancies in technologies, the marginal cell-type distributions and expression scales are usually different between ***X***_sc_ and ***X***_st_, i.e.

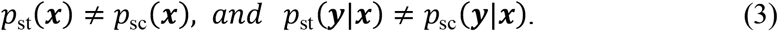

In iSORT, we employ the reference-based co-embedding approach as Seurat^37^ does. ***X***_sc_ is set as the query and ***X***_st_ is as the reference. Denote the variables after the co-embedding as ***Z***_sc_ and ***Z***_st_, respectively. Here, the function ***h*** maps the gene expression space to the latent space, represented by ***z*** = ***h***(***x***). Under the postulation that a cell’s spatial location is intrinsically related to its latent expression, we obtain

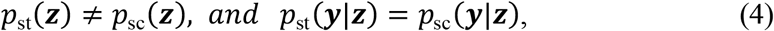

which implies that samples with similar features ***z*** have close spatial coordinates ***y***. We represent the mapping between latent expressions and locations as ***y*** = ***g***(***z***), where ***g***; ℝ*^H^* → ℝ^2^. The task is to give an estimation of ***g*** and determine the location of ***X***_sc_ by ***f***(***X***_sc_) ≔ ***g*** ∘ ***h***(***X***_sc_). Several studies have discussed the construction of ***g*** considering the covariance bias^67, 68^. iSORT addresses the estimation as a learning task to minimize a loss function

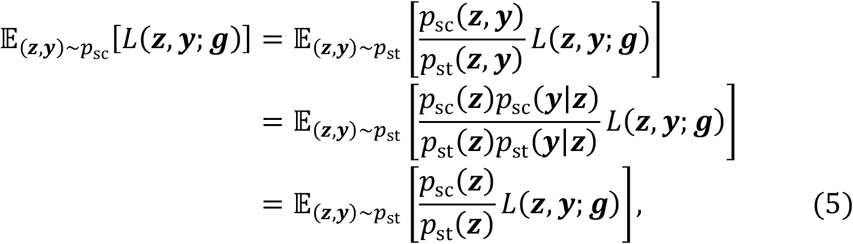

where Eq. (4) is applied in the last step, and *L*(⋅, ⋅; ***g***) is the loss function. For the spatial reconstruction, *L* is usually selected as the mean squared error loss, i.e. *L*(***z, y***; ***g***) = ‖***g***(***z***) **− *y***‖_2_. The density ratio *w*(***z***) = *p*_sc_(***z***)/*p*_st_(***z***) is the key to address the different cell-type distributions between the scRNA-seq and ST data.

For the *m* samples in the ST slice, the minimization of (5) is discretized as

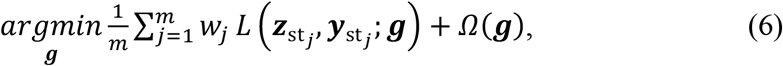

where 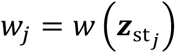 is the sample-specific density ratio and Ω(***g***) is the normalization term. Once *w***_*j*_**s are known, we can obtain ***g*** by optimizing Eq. (6) and then apply ***f*** on ***X***_sc_.

#### Estimation of the density ratio

iSORT estimates the density ratio *w*(***z***) = *p*_sc_(***z***)/*p*_st_(***z***) by the method of KLIEP^68^. KLIEP demonstrates computational efficiency, stability in performance, and effective mitigation of overfitting. Specifically, *w*(***z***) is represented linearly by a set of basis functions, i.e.

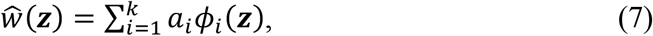

where 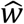 is the approximation of *w*, 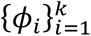 are *k* given basis functions satisfying *ϕ_i_*(***z***) ≥ 0, and *a_i_*s are the coefficients to be estimated. By default, Gaussian basis functions (GBFs) are selected as *ϕ_i_*, offering qualities of smoothness, locality, and universal approximation capability^69^.

KLIEP estimates 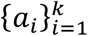 by minimizing the KL divergence between *p*_sc_(***z***) and 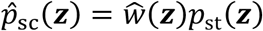, i.e.

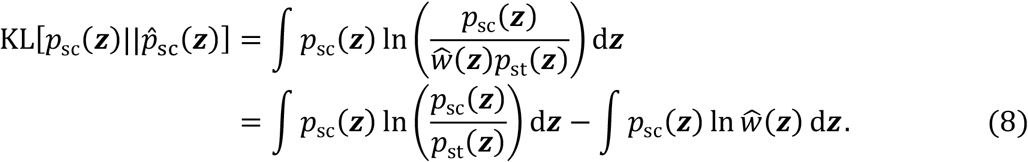

Only the second term in Eq. (8) contains *a_i_*, and the optimization is equivalent to maximize

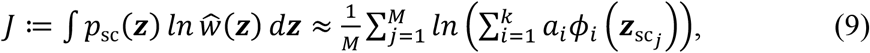

where 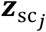 s are the latent expressions of scRNA-seq data, and *M* is the number of scRNA-seq samples. Considering the constrain of pdf as

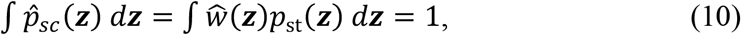

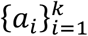 is finally estimated by solving

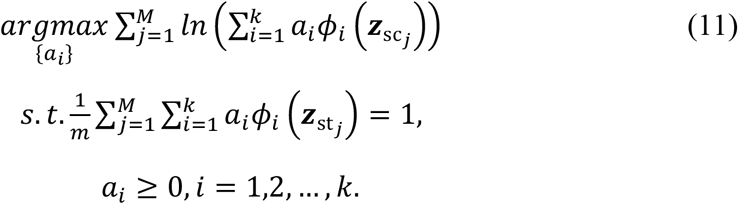

#### Optimizing the reconstruction mapping

Neural network technology can uncover the nonlinear mapping between variables with just a few hidden layers^70, 71^. To solve the optimization problem (6) and approximate spatial organization mapping ***f***, we applied a multi-layer BP neural network (Fig. 1(b)). Dropout is used to relieve the overfitting. Detailed architecture can be referred to Supplementary Note 7.

### Identification of SOGs

SOGs are the genes most relevant to the spatial organization of tissues. Within the framework of iSORT, ***f*** maps the gene expression data ***x*** ∈ ℝ*^n^* to a two-dimensional spatial coordinate ***y*** ∈ ℝ^2^, i.e. ***y*** = ***f***(***x***). We proposed the SOG index of gene *g* as

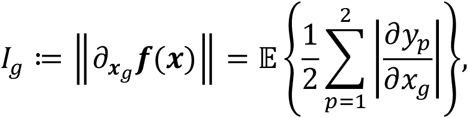

where *y*_1_, *y*_2_ are the two components of ***y, x****_g_* is the gene *g* expression in ***x***, and the expectation 𝔼 is taken over all cells/samples. SOGs are identified as the genes with top *I_g_* scores, reflecting their significant contributions to the spatial configuration and biological functionality.

### Inference of SpaRNA velocity

iSORT proposes a way named SpaRNA velocity to visualize the spatial growth of tissues. For a cell with gene expression ***x***, we suppose that its RNA velocity ***v***_RNA_ is obtained in the gene expression space by scTour^36^ or other algorithms. Using the mapping ***f***, we can define the SpaRNA velocity as

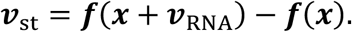

The SpaRNA velocity allows us to further explore the dynamics of tissue growth in physical space, and provides a comprehensive view of the cellular organization.

### Simulation of the coarse-grained ST data

FISH technology can provide high-resolution ST data^72^ at single-cell scale, while most sequence-based approaches such as 10X Visium can only achieve spot-resolution. The diameter of a spot in 10X Visium is 55μm, while the distance between the centers of two adjacent spots is 100μm. One spot may contain 1-10 cells.

To test the effectiveness of iSORT when using low-resolution ST data as the reference, we simulate the coarse-grained ST from the seqFISH data in mouse embryo and FISH data in *Drosophila* embryo experiments. In the simulation, we initially divided the area into a uniform grid, with the intersections of the lines serving as spots. The radius of each coarse spot is set to be one quarter of the adjacent spot spacing, and the gene expression is set as the accumulated value of all cells within the radius. We only considered spots with gene expression greater than zero.

### Collection of human artery samples and ethics statement

The human artery data were collected from patients undergoing coronary artery bypass grafting (CABG) or heart transplantation at Zhongshan Hospital, Fudan University. Written informed consent was obtained from each participant before surgery, with the study approved by the Ethics Committee of Zhongshan Hospital, Fudan University (ethical approval number: B2022-031R), and conducted in strict accordance with the principles outlined in the Declaration of Helsinki.

For scRNA-seq, artery samples were processed to generate single-cell suspensions. The process involved cell separation, labeling, and library construction, followed by sequencing on the Illumina NovaSeq platform. The sequencing data were processed with Cell Ranger (10x Genomics, version cellranger-6.0.0), aligning them to the human reference genome (GRCh38) to create a matrix of gene expression barcodes.

For ST, the samples that passed quality inspection were re-sectioned for permeabilization experiments. The Visium Spatial Tissue Optimization Slide & Reagent Kit (PN-1000193, 10X Genomics) was used to release mRNA from cells and bind it to spatially barcoded oligonucleotides on the slides. Imaging to determine the appropriate permeabilization time was performed using Leica Aperio CS2 and Leica THUNDER Imager Tissue. The libraries were then prepared using the Visium Spatial Gene Expression kit (PN-1000184, 10X Genomics).

To compare the structural and morphological differences between normal vessels and those affected by atherosclerosis, tissue sections prepared for ST were additionally stained with hematoxylin and eosin, and subsequently analyzed using a Leica Aperio CS2.

## DATA AVAILABILITY

The original public data used in this paper can be accessed through the following links: (1) FISH data from the Berkeley Drosophila Transcription Network Project (BDTNP) https://www.fruitfly.org/; (2) 10X Visium data of the human dorsolateral prefrontal cortex (DLPFC) http://spatial.libd.org/spatialLIBD/; (3) Adult mouse cortical cell datasets: GEO accession GSE71585; (4) The mouse embryo data https://marionilab.cruk.cam.ac.uk/SpatialMouseAtlas; (5) The Seurat objects ST data (10X Genomics Visium) of mouse brain https://satijalab.org/seurat/articles/spatial_vignette.html; (6)Single-cell RNA-seq data and Spatial Transcriptomics data of the developing human heart https://data.mendeley.com/datasets/mbvhhf8m62/2; (7)The human artery data: OMIX006463 on CNCB https://ngdc.cncb.ac.cn/omix/preview/SZRjY20z.

## CODE AVAILABILITY

The iSORT algorithm is available at GitHub: https://github.com/xiaojierzi/iSORT

## ACKNOWLEDGEMENTS

This work is supported by National Key R&D Program (2022YFC2704604 to J.S.), National Science Foundation of China (No. 12301620 to J.S., No. 11925103 to W.L., No. 82070463 to Y.Y.), and Science and Technology Commission of Shanghai Municipality (No. 21DZ1201402 to W.L.).

## Author contributions statement

Y.T., A.W., Q.N, and J.S. designed the research; Y.T. and J.S. conceived the method; Y.T. and J.S. conducted the numerical results; A.W. and Y.Y. conducted the experiments; Y.T., A.W., Z.W., Y.Y. and J.S. interpreted the results; W.L., Y.Y., Q.N. and J.S. supervised the work. All the authors wrote and revised the manuscript.

## REFERENCES

1. Tang, F. et al. mRNA-Seq whole-transcriptome analysis of a single cell. Nat. Methods 6, 377–382 (2009).

2. Ulrich, N. D. et al. Cellular heterogeneity of human fallopian tubes in normal and hydrosalpinx disease states identified using scRNA-seq. Dev. Cell 57, 914–929 (2022).

3. Weinreb, C., Wolock, S., Tusi, B. K., Socolovsky, M. & Klein, A. M. Fundamental limits on dynamic inference from single-cell snapshots. Proc. Natl. Acad. Sci. 115, E2467–E2476 (2018).

4. Shi, J., Li, T., Chen, L. & Aihara, K. Quantifying pluripotency landscape of cell differentiation from scRNA-seq data by continuous birth-death process. PLoS Comput. Biol. 15, e1007488 (2019).

5. Fernandez-Garcia, J. et al. CD8+ T cell metabolic rewiring defined by scRNA-seq identifies a critical role of ASNS expression dynamics in T cell differentiation. Cell Rep. 41 (2022).

6. Lambrechts, D. et al. Phenotype molding of stromal cells in the lung tumor microenvironment. Nat. Medicine 24, 1277–1289 (2018).

7. Andrews, T. S. & Hemberg, M. Identifying cell populations with scRNAseq. Mol. Aspects Medicine 59, 114–122 (2018).

8. Hu, C. et al. Cellmarker 2.0: an updated database of manually curated cell markers in human/mouse and web tools based on scrna-seq data. Nucleic Acids Res. 51, D870– D876 (2023).

9. Soneson, C. & Robinson, M. D. Bias, robustness and scalability in single-cell differential expression analysis. Nat. Methods 15, 255–261 (2018).

10. Shi, J., Teschendorff, A. E., Chen, W., Chen, L. & Li, T. Quantifying Waddington’s epigenetic landscape: a comparison of single-cell potency measures. Briefings Bioinform. 21, 248–261 (2020).

11. Ståhl, P. L. et al. Visualization and analysis of gene expression in tissue sections by spatial transcriptomics. Science 353, 78–82 (2016).

12. Moffitt, J. R. et al. Molecular, spatial, and functional single-cell profiling of the hypothalamic preoptic region. Science 362, eaau5324 (2018).

13. Eng, C.-H. L. et al. Transcriptome-scale super-resolved imaging in tissues by RNA seqFISH+. Nature 568, 235–239 (2019).

14. Wang, X. et al. Three-dimensional intact-tissue sequencing of single-cell transcriptional states. Science 361, eaat5691 (2018).

15. 10x Genomics: Visium Spatial Gene Expression. 10x Genomics https://www.10xgenomics.com/ (2024).

16. Rodriques, S. G. et al. Slide-seq: A scalable technology for measuring genome-wide expression at high spatial resolution. Science 363, 1463–1467 (2019).

17. Stickels, R. R. et al. Highly sensitive spatial transcriptomics at near-cellular resolution with Slide-seqV2. Nat. Biotechnol. 39, 313–319 (2021).

18. Chen, A. et al. Spatiotemporal transcriptomic atlas of mouse organogenesis using dna nanoball-patterned arrays. Cell 185, 1777–1792 (2022).

19. Elosua-Bayes, M., Nieto, P., Mereu, E., Gut, I. & Heyn, H. SPOTlight: seeded NMF regression to deconvolute spatial transcriptomics spots with single-cell transcriptomes. Nucleic Acids Res. 49, e50–e50 (2021).

20. Kleshchevnikov, V. et al. Cell2location maps fine-grained cell types in spatial transcriptomics. Nat. Biotechnol. 40, 661–671 (2022).

21. Biancalani, T. et al. Deep learning and alignment of spatially resolved single-cell transcriptomes with Tangram. Nat. Methods 18, 1352–1362 (2021).

22. Hao, M. et al. Stem enables mapping of single-cell and spatial transcriptomics data with transfer learning. Commun. Biol. 7, 56 (2024).

23. Nitzan, M., Karaiskos, N., Friedman, N. & Rajewsky, N. Gene expression cartography. Nature 576, 132–137 (2019).

24. Cang, Z. & Nie, Q. Inferring spatial and signaling relationships between cells from single cell transcriptomic data. Nat. Commun. 11, 2084 (2020).

25. Ren, X. et al. Reconstruction of cell spatial organization from single-cell rna sequencing data based on ligand-receptor mediated self-assembly. Cell Res. 30, 763–778 (2020).

26. Qian, J. et al. Reconstruction of the cell pseudo-space from single-cell RNA sequencing data with scSpace. Nat. Commun. 14, 2484 (2023).

27. Zhang, Q. et al. Leveraging spatial transcriptomics data to recover cell locations in single-cell RNA-seq with CeLEry. Nat. Commun. 14, 4050 (2023).

28. Wei, R. et al. Spatial charting of single-cell transcriptomes in tissues. Nat. Biotechnol. 40, 1190–1199 (2022).

29. Li, B. et al. Benchmarking spatial and single-cell transcriptomics integration methods for transcript distribution prediction and cell type deconvolution. Nat. Methods 19, 662–670 (2022).

30. Li, H. et al. A comprehensive benchmarking with practical guidelines for cellular deconvolution of spatial transcriptomics. Nat. Commun. 14, 1548 (2023).

31. Svensson, V., Teichmann, S. A. & Stegle, O. SpatialDE: identification of spatially variable genes. Nat. Methods 15, 343–346 (2018).

32. Shi, J., Chen, L. & Aihara, K. Embedding entropy: a nonlinear measure of dynamical causality. Journal of The Royal Society Interface 19, 20210766 (2022).

33. Rubin, D. B. Causal inference using potential outcomes: Design, modeling, decisions. J. Am. Stat. Assoc. 100, 322–331 (2005).

34. La Manno, G. et al. RNA velocity of single cells. Nature 560, 494–498 (2018).

35. Li, T., Shi, J., Wu, Y. & Zhou, P. On the mathematics of rna velocity i: theoretical analysis. bioRxiv 2020–09 (2020).

36. Li, Q. sctour: a deep learning architecture for robust inference and accurate prediction of cellular dynamics. Genome Biol. 24, 149 (2023).

37. Stuart, T. et al. Comprehensive integration of single-cell data. Cell 177, 1888–1902 (2019).

38. Maynard, K. R. et al. Transcriptome-scale spatial gene expression in the human dorsolateral prefrontal cortex. Nat. Neurosci. 24, 425–436 (2021).

39. Lohoff, T. et al. Integration of spatial and single-cell transcriptomic data elucidates mouse organogenesis. Nat. Biotechnol. 40, 74–85 (2022).

40. Tasic, B. et al. Adult mouse cortical cell taxonomy revealed by single cell transcriptomics. Nat. Neurosci. 19, 335–346 (2016).

41. Berkeley Drosophila Transcription Network Project. Available at: http://bdtnp.lbl.gov/.

42. Luengo Hendriks, C. L. et al. Three-dimensional morphology and gene expression in the Drosophila blastoderm at cellular resolution I: data acquisition pipeline. Genome Biol. 7, 1–21 (2006).

43. Nüsslein-Volhard, C. & Wieschaus, E. Mutations affecting segment number and polarity in Drosophila. Nature 287, 795–801 (1980).

44. Moran, P. A. Notes on continuous stochastic phenomena. Biometrika 37, 17–23 (1950).

45. Libby, P. The changing landscape of atherosclerosis. Nature 592, 524–533 (2021).

46. Sun, J. et al. Spatial transcriptional mapping reveals site-specific pathways underlying human atherosclerotic plaque rupture. J. Am. Coll. Cardiol. 81, 2213–2227 (2023).

47. Scherberich, A. et al. Tenascin-W is found in malignant mammary tumors, promotes alpha8 integrin-dependent motility and requires p38MAPK activity for BMP-2 and TNF-alpha induced expression in vitro. Oncogene 24, 1525–1532 (2005).

48. Neidhardt, J., Fehr, S., Kutsche, M., Löhler, J. & Schachner, M. Tenascin-N: characterization of a novel member of the tenascin family that mediates neurite repulsion from hippocampal explants. Mol. Cell. Neurosci. 23, 193–209 (2003).

49. Meloty-Kapella, C. V., Degen, M., Chiquet-Ehrismann, R. & Tucker, R. P. Effects of tenascin-W on osteoblasts in vitro. Cell Tissue Res. 334, 445–455 (2008).

50. Zhou, A.-X., Hartwig, J. H. & Akyürek, L. M. Filamins in cell signaling, transcription and organ development. Trends Cell Biol. 20, 113–123 (2010).

51. Seo, W. & Ziltener, H. J. CD43 processing and nuclear translocation of CD43 cytoplasmic tail are required for cell homeostasis. Blood, The J. Am. Soc. Hematol. 114, 3567–3577 (2009).

52. Meng, X.-D. et al. Knockdown of GAS5 inhibits atherosclerosis progression via reducing EZH2-mediated ABCA1 transcription in ApoE-/-mice. Mol. Ther. Acids 19, 84–96 (2020).

53. Hoerder-Suabedissen, A. & Molnár, Z. Development, evolution and pathology of neocortical subplate neurons. Nat. Rev. Neurosci. 16, 133–146 (2015).

54. Ren, H., Walker, B. L., Cang, Z. & Nie, Q. Identifying multicellular spatiotemporal organization of cells with spaceflow. Nat. Commun. 13, 4076 (2022).

55. Asp, M. et al. A spatiotemporal organ-wide gene expression and cell atlas of the developing human heart. Cell 179, 1647–1660 (2019).

56. England, M. A. The developing human: clinically oriented embryology. J. Anat. 166, 270 (1989).

57. Franco, D. & Kelly, R. G. Contemporary cardiogenesis: new insights into heart development. Cardiovasc. Res. 91, 183–184 (2011).

58. Cui, H. et al. scgpt: toward building a foundation model for single-cell multi-omics using generative ai. Nat. Methods 1–11 (2024).

59. Hao, M. et al. Large-scale foundation model on single-cell transcriptomics. Nat. Methods 1–11 (2024).

60. Efremova, M., Vento-Tormo, M., Teichmann, S. A. & Vento-Tormo, R. Cellphonedb: inferring cell–cell communication from combined expression of multi-subunit ligand–receptor complexes. Nat. Protoc. 15, 1484–1506 (2020).

61. Jin, S. et al. Inference and analysis of cell-cell communication using cellchat. Nat. Commun. 12, 1088 (2021).

62. Bergen, V., Lange, M., Peidli, S., Wolf, F. A. & Theis, F. J. Generalizing rna velocity to transient cell states through dynamical modeling. Nat. Biotechnol. 38, 1408–1414 (2020).

63. Trapnell, C. et al. The dynamics and regulators of cell fate decisions are revealed by pseudotemporal ordering of single cells. Nat. Biotechnol. 32, 381–386 (2014).

64. Street, K. et al. Slingshot: cell lineage and pseudotime inference for single-cell transcriptomics. BMC Genomics 19, 1–16 (2018).

65. Wang, G. et al. Construction of a 3d whole organism spatial atlas by joint modelling of multiple slices with deep neural networks. Nat. Mach. Intell. 5, 1200–1213 (2023).

66. Wolf, F. A., Angerer, P. & Theis, F. J. SCANPY: large-scale single-cell gene expression data analysis. Genome Biol. 19, 1–5 (2018).

67. Huang, J., Gretton, A., Borgwardt, K., Schölkopf, B. & Smola, A. Correcting sample selection bias by unlabeled data. Adv. Neural Inf. Process. Syst. 19 (2006).

68. Sugiyama, M. et al. Direct importance estimation for covariate shift adaptation. Ann. Inst. Stat. Math. 60, 699–746 (2008).

69. Poggio, T. & Girosi, F. Networks for approximation and learning. Proc. IEEE 78, 1481–1497 (1990).

70. LeCun, Y., Bengio, Y. & Hinton, G. Deep learning. Nature 521, 436–444 (2015).

71. Hornik, K., Stinchcombe, M. & White, H. Multilayer feedforward networks are universal approximators. Neural Netw. 2, 359–366 (1989).

72. Bressan, D., Battistoni, G. & Hannon, G. J. The dawn of spatial omics. Science 381, eabq4964 (2023).

