## Supplemental Information for "Transfer learning of multicellular organization via single-cell and spatial transcriptomics"

### 1 Supplementary notes

#### 1.1 Supplementary Note 1: iSORT for the multi-slice case.

##### Spatial organization mapping with multiple slices

For the multi-slice case, we have scRNA-seq with $l$ ST slices as references (See Fig. 1(b) in the main text). Denote $\boldsymbol{X}_{\text{sc}} = \{\boldsymbol{x}_{\text{sc}_1}, \boldsymbol{x}_{\text{sc}_2}, \ldots, \boldsymbol{x}_{\text{sc}_M}\}$ as the scRNA-seq expression, $\boldsymbol{X}_{\text{st}}^{(i)} = \{\boldsymbol{x}_{\text{st}_1}^{(i)}, \boldsymbol{x}_{\text{st}_2}^{(i)}, \ldots, \boldsymbol{x}_{\text{st}_{m_i}}^{(i)}\}$ as the expression of the $i$th ST slice and $\boldsymbol{Y}_{\text{st}}^{(i)} = \{\boldsymbol{y}_{\text{st}_1}^{(i)}, \boldsymbol{y}_{\text{st}_2}^{(i)}, \ldots, \boldsymbol{y}_{\text{st}_{m_i}}^{(i)}\}$ as the spatial coordinates of the $i$th ST slice, where $i = 1, 2, \ldots, l$ is the number of ST slices, and $M$ and $m_i$ are the sample sizes. The latent expressions after the co-embedding step are $\boldsymbol{Z}_{\text{sc}} = \{\boldsymbol{z}_{\text{sc}_1}, \boldsymbol{z}_{\text{sc}_2}, \ldots, \boldsymbol{z}_{\text{sc}_M}\}$ and $\boldsymbol{Z}_{\text{st}}^{(i)} = \{\boldsymbol{z}_{\text{st}_1}^{(i)}, \boldsymbol{z}_{\text{st}_2}^{(i)}, \ldots, \boldsymbol{z}_{\text{st}_{m_i}}^{(i)}\}$, respectively. The challenge is that the scRNA-seq and ST slices are usually sampled from different experiments with different tissue shapes and cell-type distributions, i.e.

$$p_{\text{sc}}(\boldsymbol{z}) \neq p_{\text{st}^{(1)}}(\boldsymbol{z}) \neq p_{\text{st}^{(2)}}(\boldsymbol{z}) \neq \ldots \neq p_{\text{st}^{(l)}}(\boldsymbol{z}), \tag{1}$$

and

$$p_{\text{st}^{(1)}}(\boldsymbol{y}|\boldsymbol{z}) \neq p_{\text{st}^{(2)}}(\boldsymbol{y}|\boldsymbol{z}) \neq \ldots \neq p_{\text{st}^{(l)}}(\boldsymbol{y}|\boldsymbol{z}), \tag{2}$$

where $p_{\text{sc}}$ and $p_{\text{st}^{(i)}}$ are the corresponding pdfs. To address this issue, iSORT generalizes the single-slice case to the minimization problem as [1] does:

$$\underset{\boldsymbol{f}}{\operatorname{argmin}} \sum_{i=1}^{l} \frac{\beta_i}{m_i} \sum_{j=1}^{m_i} w_j^{(i)} L(\boldsymbol{z}_{\text{st}_j}^{(i)}, \boldsymbol{y}_{\text{st}_j}^{(i)}; \boldsymbol{f}) + \Omega(\boldsymbol{f}), \tag{3}$$

where $\beta_i$ is the slice-specific weight integrating the heterogeneity of different slices, and $w_j^{(i)}$ is the sample-specific weight calculated by density ratio. Similar to the single-slice case, $L$ is set as the mean squared error loss, and $w_j^{(i)}$ is estimated by the KLIEP. Once $\beta_i$s are obtained, the mapping $\boldsymbol{f}$ can be approximated by a multi-layer BP neural network (Fig. 1(b) in the main text). The pseudo-space estimation is given by $\boldsymbol{f}(\boldsymbol{Z}_{\text{sc}})$. When $l = 1$, the procedure naturally degenerates to the single-slice case.

##### Estimation of the slice-specific weight

The slice-specific weight $\boldsymbol{\beta} = (\beta_1, \beta_2, \ldots, \beta_l)^T$ is used to balance the importance of multiple slices. We estimate $\boldsymbol{\beta}$ similarly as [1] does by

$$\underset{\boldsymbol{\beta}:\boldsymbol{\beta}^T\mathbf{1}=1,\ \boldsymbol{\beta}\geq 0}{\operatorname{argmin}} \sum_{i,j=1}^{M} \rho_{ij} ||\boldsymbol{\beta}^T \boldsymbol{Y}_{\text{sc}}^{(i)} - \boldsymbol{\beta}^T \boldsymbol{Y}_{\text{sc}}^{(j)}||_2^2, \tag{4}$$

where $\boldsymbol{Y}_{\text{sc}}^{(i)} = (\boldsymbol{f}^{(1)}(\boldsymbol{z}_{\text{sc}_i}), \boldsymbol{f}^{(2)}(\boldsymbol{z}_{\text{sc}_i}), \ldots, \boldsymbol{f}^{(l)}(\boldsymbol{z}_{\text{sc}_i}))^T$, $\boldsymbol{f}^{(i)}$ is the single-slice mapping estimated by scRNA-seq and the $i$th ST slice, and $\rho_{ij}$ is the value of the correlation coefficient between $\boldsymbol{z}_{\mathrm{sc}_i}$ and $\boldsymbol{z}_{\mathrm{sc}_j}$. Eq. (4) implies that with the help of $\boldsymbol{\beta}$, the predictions using different slices between cells with high similarity become smaller and the differences between cells with low similarity become larger.

Eq. (4) is a convex quadratic optimization problem with an equivalent form:

$$\underset{\boldsymbol{\beta}:\boldsymbol{\beta}^T\mathbf{1}=1,\ \boldsymbol{\beta}\geq 0}{\operatorname{argmin}} \quad \sum_{k=1}^{2} \boldsymbol{\beta}^T \hat{\boldsymbol{Y}}_{\mathrm{sc}}^{[k]} \boldsymbol{L} (\hat{\boldsymbol{Y}}_{\mathrm{sc}}^{[k]})^T \boldsymbol{\beta}, \tag{5}$$

where $\hat{\boldsymbol{Y}}_{\mathrm{sc}}^{[k]} = (\boldsymbol{Y}_{\mathrm{sc}}^{(1)}(k), \boldsymbol{Y}_{\mathrm{sc}}^{(2)}(k), \ldots, \boldsymbol{Y}_{\mathrm{sc}}^{(M)}(k))$ is the axis-$k$ of the single-slice mappings, and $\boldsymbol{L}$ is a $M \times M$ matrix with

$$\boldsymbol{L}_{ij} = \begin{cases} -\rho_{ij}, & i \neq j, \\ \sum_{k=1}^{M} \rho_{ik}, & i = j. \end{cases} \tag{6}$$

#### 1.2 Supplementary Note 2: Description of other integrative methods.

- **scSpace** scSpace[2] is designed to integrate single-cell RNA sequencing (scRNA-seq) and spatial transcriptomics (ST) data for predicting the pseudo-space of scRNA-seq. It involves initially applying transfer component analysis to merge scRNA-seq and ST data into a unified latent space, thereby extracting shared characteristics. Subsequently, a multi-layer perceptron model is trained, using the latent features as inputs and spatial coordinates as the output. This trained model is then employed to reconstruct the spatial coordinates for scRNA-seq from the single-cell data matrix. Using the gene expression profiles and pseudo-spatial information, scSpace then identifies spatially variable cell subpopulations within the scRNA-seq data.
- **Tangram** Tangram[3] is designed for spot deconvolution. It is a computational method to align scRNA-seq data with spatially resolved transcriptomic datasets. This approach maps the comprehensive scRNA-seq profiles onto the spatial domains represented in ST. Tangram operates by adapting a machine learning model to understand the distribution of gene expression across different spatial regions. The outcome is a high-resolution map of cell types and states, spatially organized according to the transcriptomic data. Since Tangram uses a probabilistic model and thus naturally predicts the spatial location of scRNA-seq. In our benchmark experimental comparisons, we used maximum likelihood estimation to make inferences about the spatial location of scRNA-seq, but this was not the only method.
- **novoSpaRc** novoSpaRc[4] is designed to impute the undetected genes. The main goal of this method is to infer the spatial distribution of cells from scRNA-seq data in the absence of a priori information on spatial location. It can also integrate ST information, which enhances the accuracy of the results. With the assumption of cells in gene expression space may be closer together in physical space, novoSpaRc uses optimal transport to depict the probability of each scRNA-seq appearing at a possible target location. Therefore, it also naturally predicts the spatial location of scRNA-seq. In our benchmark experimental comparisons, we also used maximum likelihood estimation to make inferences about the spatial location of scRNA-seq.

- **CeLEry** CeLEry[5] is designed to predict the spatial position of scRNA-seq. CeLEry predicts the spatial location of scRNA-seq by integrating the spatial transcriptome and scRNA-seq. By means of data enhancement, CeLEry enhances the number of samples in the spatial transcriptome. Subsequently, through deep learning, CeLEry uses the ST data processed through data augmentation to train the model for prediction with scRNA-seq data. Depending on the task, CeLEry's final predictions can be either regions where scRNA-seq is likely to be representative or specific coordinates.
- **CellTrek** CellTrek[6] is a computational method designed for cell tracking and spatial localization. It is primarily used for analysing and interpreting scRNA-seq data, especially when considering the spatial distribution of cells in tissues. The core function of CellTrek is to combine scRNA-seq data with imaging data at high spatial resolution, in order to understand, at the single-cell level, the exact location and environment of cells in tissues. By using interpolation as well as random forests, CellTrek is able to assign spatial locations to scRNA-seq data to provide an intuitive spatial map of cell distribution in tissues.

#### 1.3 Supplementary Note 3: Description of methods of finding spatially variable genes.

- **SpatialDE** SpatialDE[7] is a method for identifying and characterizing spatially variable genes (SVGs). It uses Gaussian process regression to analyze gene expression variability, separating it into spatial and non-spatial components through two types of random effects. The first is a spatial variance term, which models the covariance of gene expression based on the pairwise distances between samples. The second is a noise term that accounts for variability not related to spatial components. This method quantifies the proportion of variance explained by these components, providing a measure of spatial variance. SpatialDE detects significant spatially variable genes (SVGs) by comparing this comprehensive model with a reduced model that excludes the spatial variance component.
- **Moran's** $I$ Moran's $I$ is a statistical measure used to determine spatial autocorrelation[8], where values range from -1 (perfect dispersion) to +1 (perfect correlation). It identifies genes whose expression levels show significant spatial patterns across different tissue samples or within a spatially structured tissue context. By applying Moran's $I$, researchers can pinpoint genes that exhibit non-random spatial distributions, aiding in the understanding of spatial heterogeneity in gene expression which can be pivotal for targeted therapeutic strategies and understanding cellular interactions in complex tissues. We use Squidpy[9] to calculate Moran's $I$ which is defined as:

$$I = \frac{n}{W} \frac{\sum_{i=1}^{n} \sum_{j=1}^{n} W_{ij} z_i z_j}{\sum_{i=1}^{n} z_i^2}$$

  where $z_i$ is the deviation of the feature from the mean $(x_i - \overline{X})$, $w_{ij}$ is the spatial weight between observations, $n$ is the number of spatial units and $W$ is the sum of all $w_{ij}$.

#### 1.4 Supplementary Note 4: Description of datasets.

- **Human dorsolateral prefontal cortex (DLPFC) dataset.** The DLPFC dataset is collected from KR Maynard et al[10]. It contains a ST dataset, which was sequenced by 10X Visium, featuring 12 slices from three independent adult donors without any neurological symptoms. Each donor has 4 slices: ID151507, ID151508, ID151509, and ID151510 (Donor 1); ID151669, ID151670, ID151671, and ID151672 (Donor 2); ID151673, ID151674, ID151675, and ID151676 (Donor 3). The dataset is well-annotated for seven spatial domains including six cortical layers and white matter. All raw and processed data, along with the code, are publicly available to the scientific community and can be accessed through the spatialLIBD web application: http://spatial.libd.org/spatialLIBD/.

  1. **Reconstruction experiment in Fig. 2(a)**: In the reconstruction experiment in Fig. 2(a), we used ST slices ID151674 and ID151675. The ST slice ID151674 was taken as the scRNA-seq input by removing its spatial coordinates. The ST slice ID151675 was used as the ST reference.
  2. **Reconstruction experiments with multi-slice references in Fig. 3(a)**: In the reconstruction experiments in Fig. 3(a), the ST slice ID151674 was taken as the scRNA-seq input by removing its spatial coordinates. In case I, we used ST slice ID151675 as ST reference. In case II, we used ST slice ID151671 as ST reference. In case II', we used ST slice ID151507 as ST reference. In case III, we used ST slices ID151673, ID151675 and ID151676 as references. In case IV, we used ST slices ID151507, ID151675, and ID151671 as references. In case IV', we used ST slices ID151675, ID151507, and ID151508 as references. In case IV", we used ST slices ID151675, ID151670, and ID151671 as references. In case V, we used ST slices ID151673, ID151675, and ID151676 rotated by 45, 0, and $-45$ degree respectively as references. In case VI, we used ST slices ID151507, ID151675 and ID151671 rotated by 30, 0, and $-40$ degree respectively as references.
  3. **In-silico gene knockout experiments in Fig. 5(b) and (c)**: In the in-silico gene knockout experiments in Fig. 2(a), we used ST slices ID151674 and ID151675. The ST slice ID151674 was taken as the scRNA-seq input by removing its spatial coordinates. The ST slice ID151675 was used as the ST reference. For the in-silico knockout experiment, we set the expression of knockout genes as zero.
  4. **Visualization of spatial velocity experiments in Figs. 6(a)-(f)**: In spatial velocity experiments, we used ST slice ID151674.

- **Mouse embryo dataset.** The mouse embryo dataset is collected from T Lohoff et al[11]. It contains a ST dataset, which used sequential fluorescence in situ hybridization (seqFISH), an image-based single-cell transcriptomics technique, to identify mRNAs in tissue sections from three mouse embryos at the 8–12 somite stage. It has three ST slices.

  **Reconstruction experiments in Fig. 2(c)**: In the reconstruction experiment in Fig. 2(c), we selected the 'embryo1' sample from the mouse embryo dataset and removed cells labeled as "NA" or "Low quality" to ensure the quality of the data for analysis. Subsequently, we extracted the first and second dimensional spatial coordinates of the cells. Using these two dimensions, we simulated a coarse-grained ST reference. The scRNA-seq input was taken as the gene expression from each spot by removing its spatial coordinates.

- **Mouse brain dataset.** The mouse brain data are collected from T Lohoff et al[12] and https://satijalab.org/seurat/articles/spatial_vignette.html, containing a scRNA-seq dataset[12] and a ST dataset, respectively. The scRNA-seq dataset was sequenced by Smart-seq2 and the ST dataset was sequenced by 10X Visium.
  **Reconstruction experiments in Fig. 2(d)**: In this experiment, the scRNA-seq and ST data are processed as the pipeline of iSORT.
- ***Drosophila* embryonic development dataset.** The *Drosophila* embryonic development dataset is collected from Berkeley Drosophila Transcription Network Project (BDTNP)[13, 14]. It has one ST slice and was sequenced by FISH.
  **Reconstruction experiments in Fig. 4**: The original data presents in three-dimensional space, and we collected the data from the x-z plane. The seven-stripe pattern of genes such as '*ftz*' is clear to be observed. In this plane, we obtained a low-resolution ST reference through coarse-graining simulation. The scRNA-seq input was obtained by removing the spatial information.
- **Human artery dataset.** The human artery dataset was collected by Zhongshan Hospital, Fudan University. It is composed of scRNA-seq data and ST data. The ST data has 8 slices.
  **Collection of human artery samples and ethics statement**
  The human artery data were collected from patients undergoing coronary artery bypass grafting (CABG) or heart transplantation at Zhongshan Hospital, Fudan University. Written informed consent was obtained from each participant before surgery, with the study approved by the Ethics Committee of Zhongshan Hospital, Fudan University (ethical approval number: B2022-031R), and conducted in strict accordance with the principles outlined in the Declaration of Helsinki.
  For scRNA-seq, artery samples were processed to generate single-cell suspensions. The process involved cell separation, labeling, and library construction, followed by sequencing on the Illumina NovaSeq platform. The sequencing data were processed with Cell Ranger (10x Genomics, version cellranger-6.0.0), aligning them to the human reference genome (GRCh38) to create a matrix of gene expression barcodes.
  For ST, the samples that passed quality inspection were re-sectioned for permeabilization experiments. The Visium Spatial Tissue Optimization Slide & Reagent Kit (PN-1000193, 10X Genomics) was used to release mRNA from cells and bind it to spatially barcoded oligonucleotides on the slides. Imaging to determine the appropriate permeabilization time was performed using Leica Aperio CS2 and Leica THUNDER Imager Tissue. The libraries were then prepared using the Visium Spatial Gene Expression kit (PN-1000184, 10X Genomics).
  **Reconstruction experiments in Fig. 5(h)**: In this reconstruction experiments, there were two independent scRNA-seq samples as inputs: one from an individual without atherosclerosis (AS), and the other from a patient with AS. Three ST slices were used as multiple heterologous references: two from individuals without AS, and one from a patient with AS. We applied a reversible polar transformation to the ST

data in preprocessing before using iSORT to reconstruct the spatial organization (see Supplementary Notes 6).

- **Human developmental heart dataset.** The human developmental heart dataset is collected from M Asp et al[15]. It is composed of scRNA-seq data and ST data. The scRNA-seq of the human developmental heart tissues was performed using the 10x Chromium platform, and the STs were processed using the Nordiag Magnatrix 8000+ automated system. Four human developmental heart tissues were used in the study. Post-conceptional and clinical age were determined using clinical ultrasound and stage-dependent anatomical landmarks of the embryos: 4.5-5, 6.5, 6.5-7, and 9 post-conception weeks (PCWs). There are 4, 9 and 6 ST slices in the 4.5-5 PCW, 6.5 PCW and 9 PCW respectively.

  **Visualization of spatial velocity experiments in Fig. 6(h)**: In this dataset, the scRNA-seq data is well-annotated, while the ST data is not. We first used Seurat to annotate the cell types in the ST data using the scRNA-seq data as a reference. In the visualization of spatial velocity experiments, we overlapped different slices from the same stage and treated them as one ST slice. As a result, we obtained three outcomes for data at 4.5-5, 6.5, and 9 PCW.

#### 1.5 Supplementary Note 5: Comparative metrics.

##### Intra-layer similarity

For a given layer/cluster $C$, the intra-layer average distance $\hat{D}_C$ is defined as

$$\hat{D}_C = \frac{1}{|C|} \sum_{\boldsymbol{x}_i \in C} \|\boldsymbol{x}_i - \boldsymbol{\mu}_C\|_2, \tag{7}$$

where $\boldsymbol{x}_i$ are samples in $C$, $\boldsymbol{\mu}_C$ is the centroid of $C$, and $|C|$ is the counts of samples. $\hat{D}_C$ measures the average distance of all points within a specific layer to the centroid. The intra-layer similarity $S_C$ is defined as

$$S_C = \psi(\hat{D}_C) := 1 - \frac{\hat{D}_C - \hat{D}_{\min}}{\hat{D}_{\max} - \hat{D}_{\min} + \epsilon} \tag{8}$$

where $\psi$ represents the normalization function, $\epsilon$ is a small constant to prevent extremes, and $\hat{D}_{\min}$ and $\hat{D}_{\max}$ are the minimum and maximum values of $\hat{D}_C$ among all intra-layer average distance of one layer by different methods, respectively. $S_C$ is between 0 and 1, and a larger value implies a better clustering result.

##### Normalized density distribution

For a layer/cluster $C$, its density distribution $\rho_C$ is defined as

$$\rho_C = \frac{\hat{\rho}_C}{\max_C \hat{\rho}_C}, \quad \text{and} \quad \hat{\rho}_C = \frac{|C|}{V(C)}, \tag{9}$$

where $|C|$ is the number of samples in $C$, and $V(C)$ is the volume of the convex hull occupied by $C$. $\rho_C$ is between 0 and 1, and evaluates the spatial concentration of data points within a given layer or cluster $C$. A larger value of $\rho_C$ indicates a better clustering.

##### Aggregative volume index and aggregative perimeter index

For a layer/cluster $C$, aggregative volume index $A_C$ and aggregative perimeter index $P_C$ are defined as

$$A_C = \psi(V_C), \quad P_C = \psi(S_C), \tag{10}$$

Where $V(C)$ is the volume of the convex hull of $C$, $S_C$ is the perimeter, and $\psi$ is the normalization function used in Eq. (8). $A_C$ and $P_C$ are between 0 and 1. Generally, larger values of $A_C$ and $P_C$ (smaller values of $V_C$ and $S_C$) indicate a more compact and well-defined layer.

##### Accuracy

We assume a selection of two-dimensional random variables that follow a uniform distribution, with their values ranging within the $x$ and $y$ axes of the actual ST data. For $n$ spots/cells in the spatial transcriptome, we generate $n$ such two-dimensional random variables. By calculating the Mean Squared Error (MSE) between these random points and the actual spatial transcriptome, we can define precision as:

$$\text{Accuracy} = \frac{\text{MSE}_{\text{random}} - \text{MSE}_{\text{reconstructed}}}{\text{MSE}_{\text{random}}}$$

where $\text{MSE}_{\text{random}}$ represents the MSE between the random points and the actual spatial transcriptome and $\text{MSE}_{\text{reconstructed}}$ represents the MSE between the reconstructed result and the actual spatial transcriptome.

#### 1.6 Supplementary Note 6: Preprocessing of spatial coordinates of artery data.

To adapt to the complex topology of vascular structures in our computational analysis, we transformed the spatial coordinates of the ST data into polar coordinates. This transformation is performed by converting Cartesian coordinates $(x, y)$ into polar coordinates $(r, \theta)$ using the following formulas:

$$r = \sqrt{x^2 + y^2} \tag{11}$$

$$\theta = \arctan(y/x) \tag{12}$$

where $r$ is the radial distance and $\theta$ is the angle in radians. This approach helps align the data with the inherent cylindrical symmetry of arteries. After the spatial reconstruction process, we applied an inverse transformation to revert the coordinates to their original Cartesian form, using these formulas:

$$x = r \cdot \cos(\theta) \tag{13}$$

$$y = r \cdot \sin(\theta) \tag{14}$$

thereby preserving the topological integrity of the data for subsequent analysis.

#### 1.7 Supplementary Note 7: Neural network architecture.

We approximate $\boldsymbol{f}$ by a multi-layer BP neural network. The architecture is described as follows:

- **Network Architecture:** The default setting includes five fully connected layers. Each hidden layer is followed by a dropout layer to prevent overfitting. Users can adjust the depth and width of the network according to the complexity of the input data.
- **Input Layer:** The input layer contains $H$ neurons, which is the number of genes. Latent expressions in $\boldsymbol{Z}_{\text{sc}}$ or $\boldsymbol{Z}_{\text{st}}$ are used as inputs.
- **Hidden Layers:** The model contains several hidden layers, and ReLU activation function is used in each layer to enhance the network's non-linear mapping capability.
- **Dropout Layer:** Dropout layers are inserted after each hidden layer with a dropout rate of 0.1 during the training process.
- **Output Layer:** The output layer has a dimension of 2, representing the spatial coordinates on the $x$-axis and $y$-axis.
- **Loss Function:** Typically, the mean squared error loss is chosen and the training process employs the Adam optimizer with a learning rate of 0.001.

#### 1.8 Supplementary Note 8: Detailed GO enrichment analysis on the top-50 SOGs in the human artery experiment.

After the reconstruction of the topology of arteries, we further explored the SOGs (supplementary Fig. 10). Based on the index $I_g$, we ranked the genes and picked out the top 20 SOGs in both cases (supplementary tables 5 and 6). In the ranking plot of gene scores, we found that both had 9 overlapping genes in the top 10 and 16 overlapping genes in the top 20, which is closely related to the fact that we used both normal and atherosclerotic arteries as references. Nonetheless, the difference in the ordering of their scores, as well as the genes specific to each of them, reflects the variability of the sample and the heterogeneity originating from the data. Among these SOGs, *TNN* achieved the highest score. *TNN*, also known as *tenascin-W*, is implicated in facilitating integrin binding, a quintessential process for cell adhesion and migration[16]. Its association with cell proliferation and potential roles in neuronal generation and osteoblast differentiation highlight its importance in vascular structure and function[17–19]. Meanwhile, a number of genes thought to be strongly associated with atherosclerosis such as *RFLNA*, *SPN*, and *EZH2* achieved high scores. *RFLNA* is hypothesized to promote filamin binding, playing a vital role in organizing actin filaments, bone mineralization, chondrocyte development, and cell motility[20–22]. Such processes are integral to the dynamic nature of vascular cells, which require structural pliability, potentially linking to the atherogenic process. Recent studies propose that *filaminA* expression could emerge as a significant prognostic biomarker for atherosclerosis[23]. *SPN*, widely recognized as *CD43*, is involved in numerous signaling pathways that enhance cell adhesion and migration[24–26]. Notably, it has been shown that anti-*CD43* can significantly reduce monocyte adhesion to endothelial cells during inflammatory responses[27], further corroborating the role of *SPN* in atherosclerotic processes. In both cases, *EZH2* was identified as SOGs. The literatures indicate that *EZH2* is highly conserved from *Drosophila* to primates and is primarily associated with gene transcriptional repression. *EZH2* is indispensable for regular cellular development, differentiation, maintenance of gene expression, and determination of cell fate[28–30]. Furthermore, its association with cardiovascular diseases, particularly atherosclerosis[31, 32], posits a potential role in vascular health maintenance. The concurrent presence of *EZH2* in both conditions may suggest a dual functionality, contributing to vascular homeostasis and pathology. This allows for a comparative observation of *EZH2*'s potential bifunctional role in the maintenance of vascular function and in disease manifestation, possibly providing fruitful avenues for future research into vascular disease biomarkers and therapeutic targets.

We then performed GO enrichment analysis on the top 50 genes ranked by SOG scores in our study. The GO analysis reveals that 11 out of the top 20 results in GO are common to both the AS and normal samples (see supplementary table 5, supplementary table 6, supplementary Fig. 11 and supplementary Fig. 12). These shared processes include the regulation of leukocyte proliferation (GO:0070663, GO:0070661), lymphocyte activation (GO:0046631, GO:0002291), and lymphocyte immune regulation (GO:0032944), among others. The presence of these shared processes suggests a commonality in core biological functions between the AS and normal samples. It is noteworthy that unique biological process terms appear in the AS samples, including but not limited to calcification and bone formation (GO: 0060348), and phosphoric diester hydrolase activity (GO: 0008081). These terms in AS samples suggest their potential involvement in specific biological processes or functions associated with abnormal mineral deposition in atherosclerosis. This emphasizes AS's capacity to affect particular biological processes related to tissue, region, or cell type, particularly those correlated with abnormal mineralization or calcification processes.

### 2 Supplementary tables

#### 2.1 Supplementary Table 1: Accuracy of different cases in reconstructing human DLPFC slice ID151674 using multiple ST references.

**Table 1** Accuracy of different cases in reconstructing human DLPFC slice ID151674 using multiple ST references

| Table of accuracy | | | | | | | | | |
|---|---|---|---|---|---|---|---|---|---|
| Case | I | II | II' | III | IV | IV' | IV" | V | VI |
| Accuracy | 74.1% | 75.4% | 68.0% | 86.2% | 82.6% | 79.7% | 81.5% | 82.6% | 79.1% |

#### 2.2 Supplementary Table 2: List of top 20 SOGs found by iSORT in DLPFC experiment.

**Table 2** List of top 20 SOGs found by iSORT in DLPFC experiment

| Gene ID | | | |
|---|---|---|---|
| ENSG00000110484 | ENSG00000153002 | ENSG00000124935 | ENSG00000096006 |
| ENSG00000173432 | ENSG00000160182 | ENSG00000178372 | ENSG00000159763 |
| ENSG00000122585 | ENSG00000170323 | ENSG00000164600 | ENSG00000111341 |
| ENSG00000175356 | ENSG00000244734 | ENSG00000251562 | ENSG00000162551 |
| ENSG00000211897 | ENSG00000048392 | ENSG00000181195 | ENSG00000196616 |

#### 2.3 Supplementary Table 3: List of top 20 SVGs found by SpatialDE in DLPFC experiment.

**Table 3** List of top 20 SVGs found by SpatialDE in DLPFC experiment

| Gene ID | | | |
|---|---|---|---|
| ENSG00000147324 | ENSG00000144230 | ENSG00000258279 | ENSG00000232229 |
| ENSG00000255366 | ENSG00000172724 | ENSG00000151413 | ENSG00000164946 |
| ENSG00000085449 | ENSG00000095539 | ENSG00000163617 | ENSG00000170473 |
| ENSG00000150455 | ENSG00000081041 | ENSG00000224109 | ENSG00000139971 |
| ENSG00000162614 | ENSG00000135334 | ENSG00000157388 | ENSG00000134531 |

#### 2.4 Supplementary Table 4: List of top 20 genes ranked by Moran's $I$ in DLPFC experiment.

**Table 4** List of top 20 Genes ranked by Moran's $I$ in DLPFC experiment

| Gene ID | | | |
|---|---|---|---|
| ENSG00000110484 | ENSG00000123560 | ENSG00000168314 | ENSG00000131095 |
| ENSG00000091513 | ENSG00000109846 | ENSG00000173786 | ENSG00000124935 |
| ENSG00000105695 | ENSG00000167641 | ENSG00000080822 | ENSG00000136541 |
| ENSG00000013297 | ENSG00000118785 | ENSG00000173432 | ENSG00000172508 |
| ENSG00000160307 | ENSG00000204655 | ENSG00000132639 | ENSG00000154146 |

#### 2.5 Supplementary Table 5: List of top 20 SOGs in diseased artery with atherosclerosis.

**Table 5** List of top 20 SOGs in diseased artery with atherosclerosis (Genes with underlined expression overlap with Supplementary Table 6)

| Gene Name | | | | |
|---|---|---|---|---|
| *TNN* | *SPN* | *SUSD4* | *TBX1* | *EZH2* |
| *PDE1C* | *HHIP* | *HELLPAR* | *RFLNA* | *MMP9* |
| *EBI3* | *FJX1* | *CSF2RB* | *PCDH20* | *FAM180B* |
| *CD72* | *HAVCR2* | *ASGR1* | *CHRM3* | *NKAPL* |

#### 2.6 Supplementary Table 6: List of top 20 SOGs in normal artery without atherosclerosis.

**Table 6** List of top 20 SOGs in normal artery without atherosclerosis (Genes with underlined expression overlap with Supplementary Table 5)

| Gene Name | | | | |
|---|---|---|---|---|
| *TNN* | *SUSD4* | *SPN* | *EZH2* | *TBX1* |
| *RFLNA* | *HHIP* | *PDE1C* | *HELLPAR* | *EBI3* |
| *PCDH20* | *CSF2RB* | *ASGR1* | *NTS* | *CD72* |
| *NKAPL* | *FJX1* | *SKAP1* | *SMPDL3A* | *LINC02269* |

#### 2.7 Supplementary Table 7: Top 20 GO terms and their descriptions in analysis of diseased artery with atherosclerosis.

**Table 7** Top 20 GO terms and their descriptions in analysis of diseased artery with atherosclerosis

| ID | Description |
|---|---|
| GO:0009897 | external side of plasma membrane |
| GO:0046631 | alpha-beta T cell activation |
| GO:0002460 | adaptive immune response based on somatic recombination of immune receptors built from immunoglobulin superfamily domains |
| GO:0008081 | phosphoric diester hydrolase activity |
| GO:0048705 | skeletal system morphogenesis |
| GO:0070663 | regulation of leukocyte proliferation |
| GO:0140375 | immune receptor activity |
| GO:0002819 | regulation of adaptive immune response |
| GO:0022407 | regulation of cell-cell adhesion |
| GO:0032944 | regulation of mononuclear cell proliferation |
| GO:1903037 | regulation of leukocyte cell-cell adhesion |
| GO:0050863 | regulation of T cell activation |
| GO:0050670 | regulation of lymphocyte proliferation |
| GO:0002291 | T cell activation via T cell receptor contact with antigen bound to MHC molecule on antigen presenting cell |
| GO:0002286 | T cell activation involved in immune response |
| GO:0042088 | T-helper 1 type immune response |
| GO:0070661 | leukocyte proliferation |
| GO:0046634 | regulation of alpha-beta T cell activation |
| GO:0060348 | bone development |
| GO:0002707 | negative regulation of lymphocyte mediated immunity |

#### 2.8 Supplementary Table 8: Top 20 GO terms and their descriptions in analysis of normal artery.

**Table 8** Top 20 GO terms and their descriptions in analysis of normal artery

| ID | Description |
|---|---|
| GO:0070663 | regulation of leukocyte proliferation |
| GO:0032640 | tumor necrosis factor production |
| GO:0032680 | regulation of tumor necrosis factor production |
| GO:0071706 | tumor necrosis factor superfamily cytokine production |
| GO:1903555 | regulation of tumor necrosis factor superfamily cytokine production |
| GO:0002460 | adaptive immune response based on somatic recombination of immune receptors built from immunoglobulin superfamily domains |
| GO:0046631 | alpha-beta T cell activation |
| GO:0070661 | leukocyte proliferation |
| GO:0042088 | T-helper 1 type immune response |
| GO:0022407 | regulation of cell-cell adhesion |
| GO:0048705 | skeletal system morphogenesis |
| GO:0002291 | T cell activation via T cell receptor contact with antigen bound to MHC molecule on antigen presenting cell |
| GO:0050670 | regulation of lymphocyte proliferation |
| GO:1903037 | regulation of leukocyte cell-cell adhesion |
| GO:0032944 | regulation of mononuclear cell proliferation |
| GO:0002707 | negative regulation of lymphocyte mediated immunity |
| GO:0002823 | negative regulation of adaptive immune response based on somatic recombination of immune receptors built from immunoglobulin superfamily domains |
| GO:0002820 | negative regulation of adaptive immune response |
| GO:0007159 | leukocyte cell-cell adhesion |
| GO:1903039 | positive regulation of leukocyte cell-cell adhesion |

### 3 Supplementary figures

#### 3.1 Supplementary Fig. 1: Heatmap of four clustering metrics.

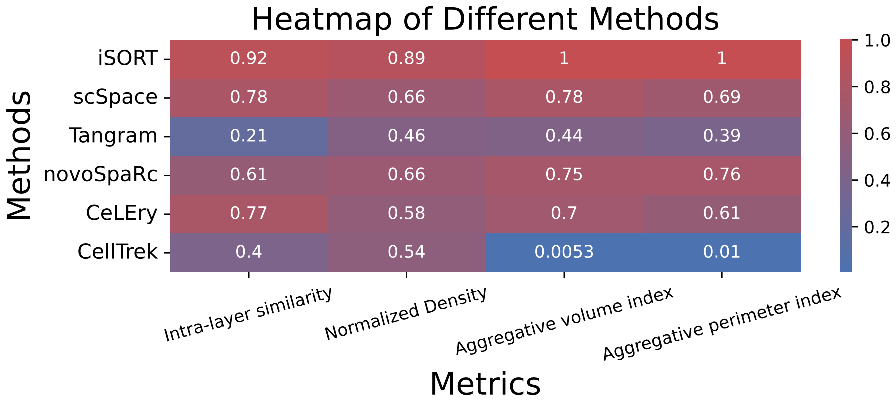

**Fig. 1 Heatmap of four clustering metrics.** Heatmap of four metrics (Mean of intra-layer similarity, mean of normalized density, mean of aggregative volume index and mean of aggregative perimeter index) of different methods (iSORT, scSpace, Tangram, novoSpaRc, CeLEry and CellTrek). The redder the colour, the better the clustering of the hierarchies. iSORT outperforms other methods in layering performance in an average sense.

##### 3.2 Supplementary Fig. 2: Comparison of different methods on mouse embyro.

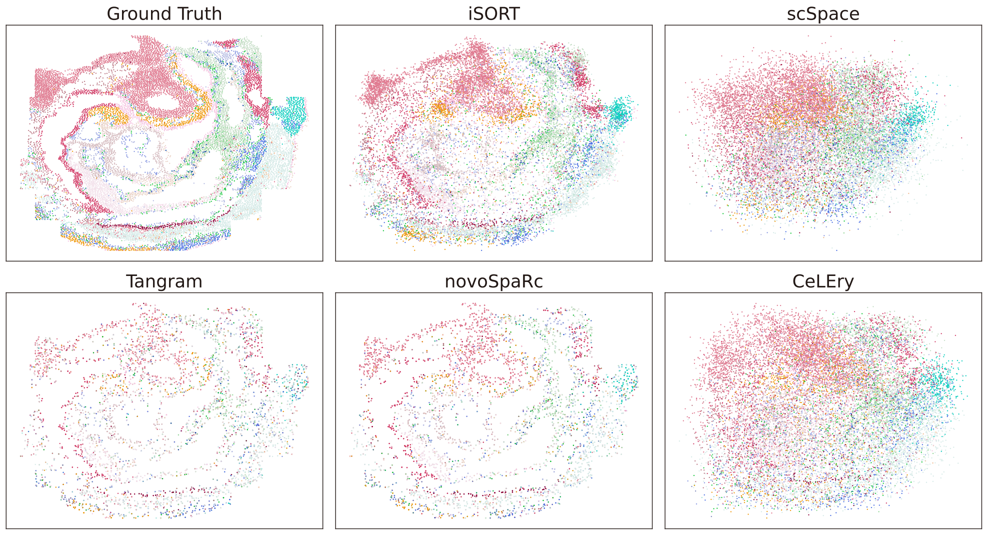

**Fig. 2 Comparison of different methods on mouse embyro.** The first row is the ground truth of mouse embyro, the reconstruction result of iSORT and the reconstruction result of scSpace from left to right. The second row is the reconstruction result of Tangram, novoSpaRc and CeLEry from left to right.

#### 3.3 Supplementary Fig. 3: Comparison of different methods on mouse brain.

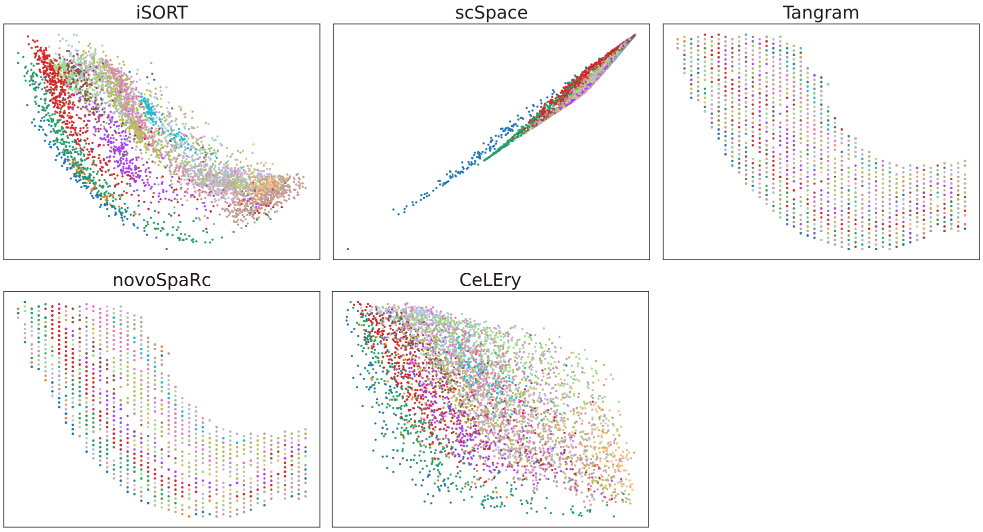

**Fig. 3 Comparison of different methods on mouse brain.** The first row is the reconstruction result of iSORT, scSpace and Tangram from left to right. The second row is the reconstruction result of novoSpaRc and CeLEry from left to right. Tangram and novoSpaRc were able to distinguish different cerebral cortices of the mouse brain well on the simulated spot space, and iSORT was able to distinguish cerebral cortices well on continuous space while scSpace and CeLEry failed.

#### 3.4 Supplementary Fig. 4: Original space of ST slices used as references in DLPFC experiments.

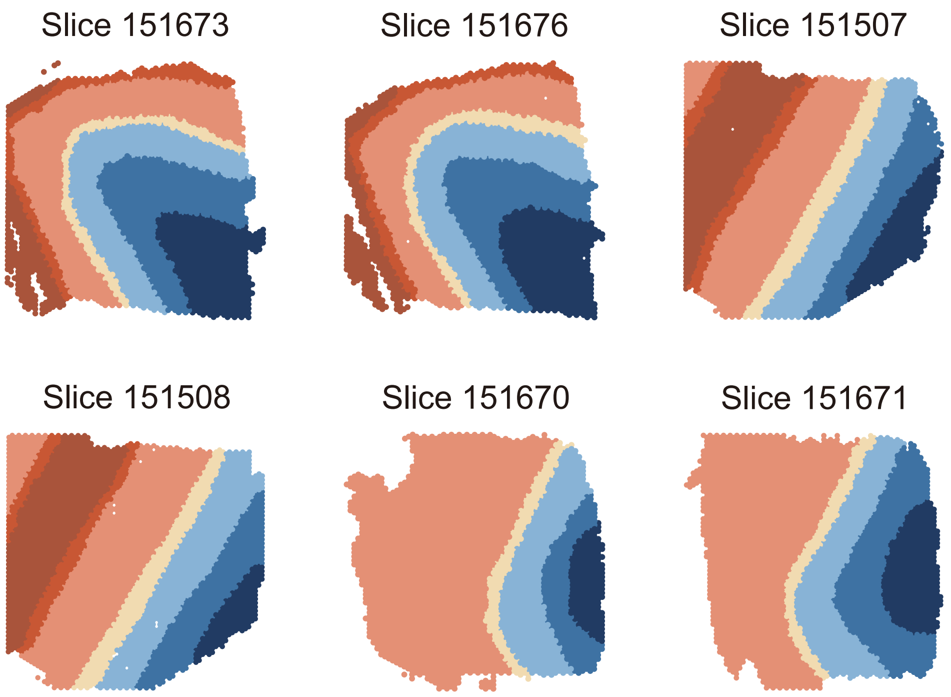

**Fig. 4 Original space of ST slices used as references in DLPFC experiments.** The first row is ID151673 (left), ID151676 (middle) and ID151507 (right) from left to right. The second row is ID151508 (left), ID151670 (middle) and ID151671 (right) from left to right. Different colors represent different cerebral cortexes.

#### 3.5 Supplementary Fig. 5: Reconstruction results of ST slice ID151674 in case II', IV' and IV".

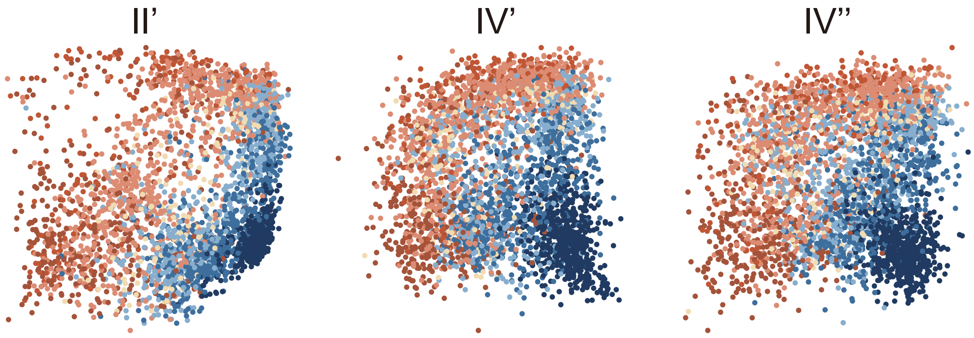

**Fig. 5 Reconstruction results of ST slice ID151674 in case II', IV' and IV".** The reference of II' is ID151507 (heterologous ST reference), the references of IV' are ID151675, ID151507, and ID151508 (heterologous ST references), the references of IV" are ID151675, ID151670, and ID151671 (heterologous ST references).

##### 3.6 Supplementary Fig. 6: Rotated DLPFC slices.

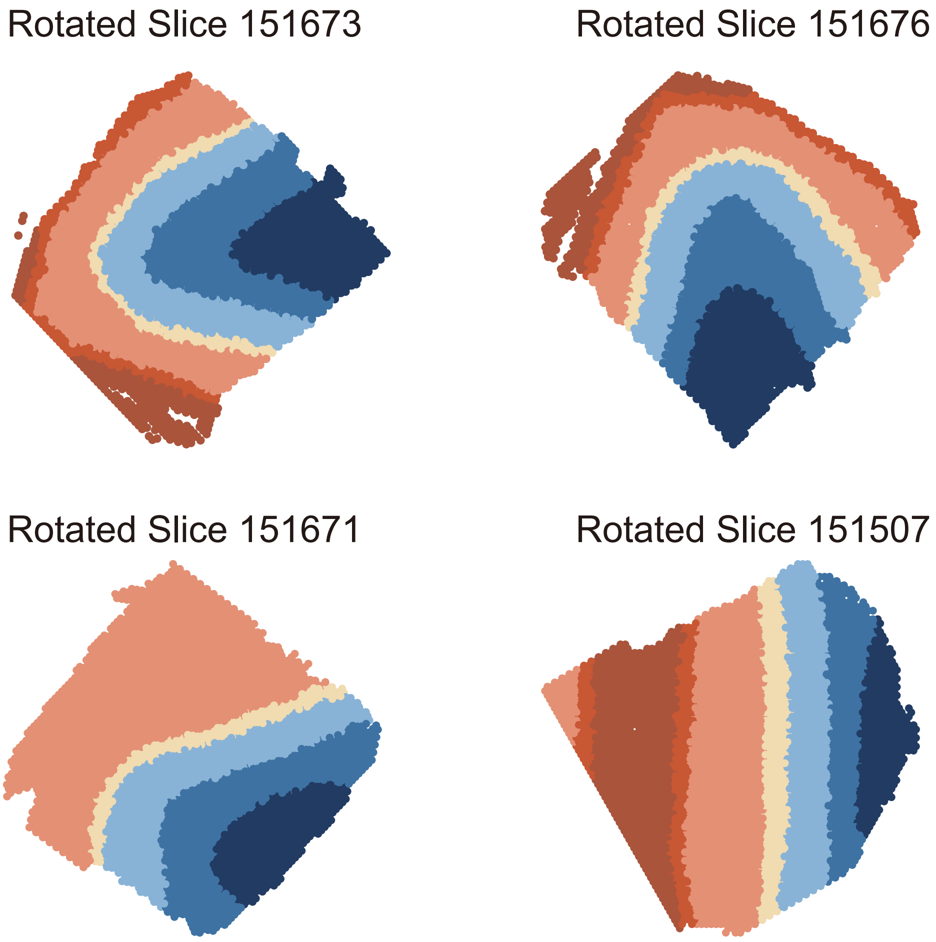

**Fig. 6 Rotated DLPFC slices.** The first row is the rotated ID151673 (45°) and ID151676 (-45°) from left to right. The second row is the rotated slice ID151671 (-40°) and ID151507 (30°) from left to right. By introducing distortion, different ST slices exhibit spatial variability.

#### 3.7 Supplementary Fig. 7: Ground truth spatial expression patterns of specific genes in *Drosophila* embyro experiment.

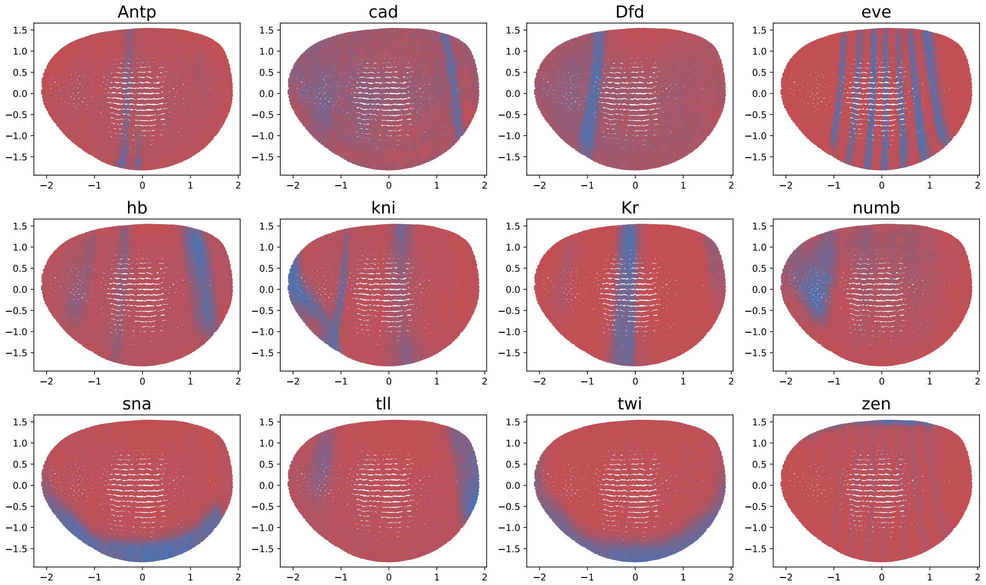

**Fig. 7 Ground truth spatial expression patterns of specific genes in *Drosophila* embyro experiment.** The following genes play a important role on *Drosophila* embyro development. *Antp* (*Antennapedia*): A homeotic gene playing a crucial role in determining the specific identities of body segments. *cad* (*caudal*): Vital for the formation of posterior structures in the embryo. Dfd (Deformed): Another homeotic gene, influencing the formation of the head and anterior thorax. *eve* (*even-skipped*): A pair-rule gene, essential for the segmentation of the body. *hb* (*hunchback*): A gap gene, critical for early embryonic segmentation and the establishment of the body axis. *kni* (*knirps*): An important gap gene. *Kr* (*Krüppel*): A gap gene, crucial for early embryonic segmentation. *numb*: Plays a key role in neural development and cell fate determination. *sna* (*snail*): Important for cell migration and morphogenesis during embryonic development. *tll* (*tailless*): A significant gap gene, affecting the development of both ends of the embryo. *twi* (*twist*): Very important for muscle development and the establishment of early embryonic morphology. *zen* (*zeno*): Crucial for the development of the dorsal part of the embryo.

#### 3.8 Supplementary Fig. 8: Simulated coarse-grained spot spatial expression patterns of specific genes in *Drosophila* embryo experiment.

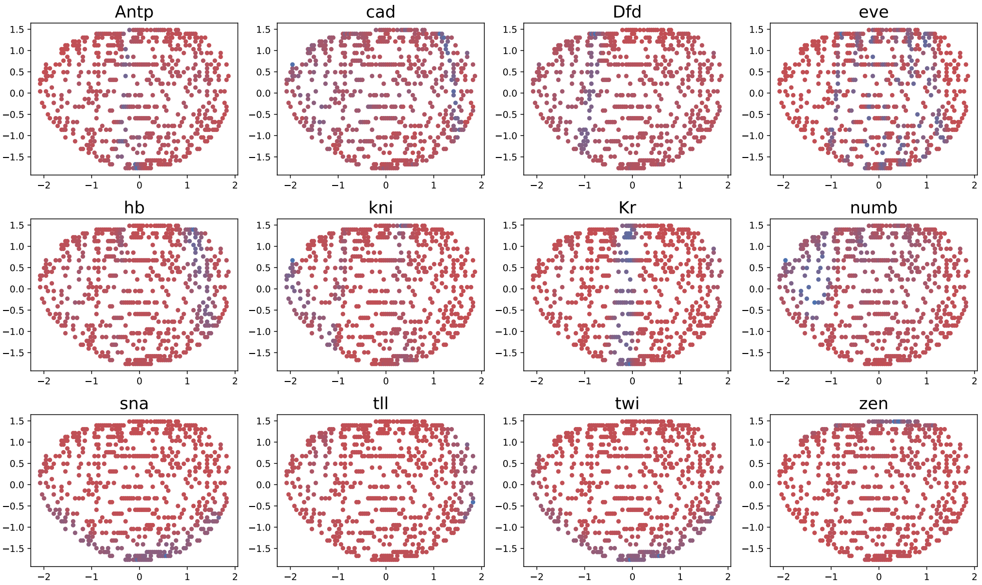

**Fig. 8 Simulated coarse-grained spots spatial expression patterns of specific genes in *Drosophila* embyro experiment.** On the simulated coarse-grained spot space, the specific spatial pattern of most of the genes could not be observed.

#### 3.9 Supplementary Fig. 9: Reconstruction results of spatial expression patterns of specific genes in *Drosophila* embyro experiment by iSORT.

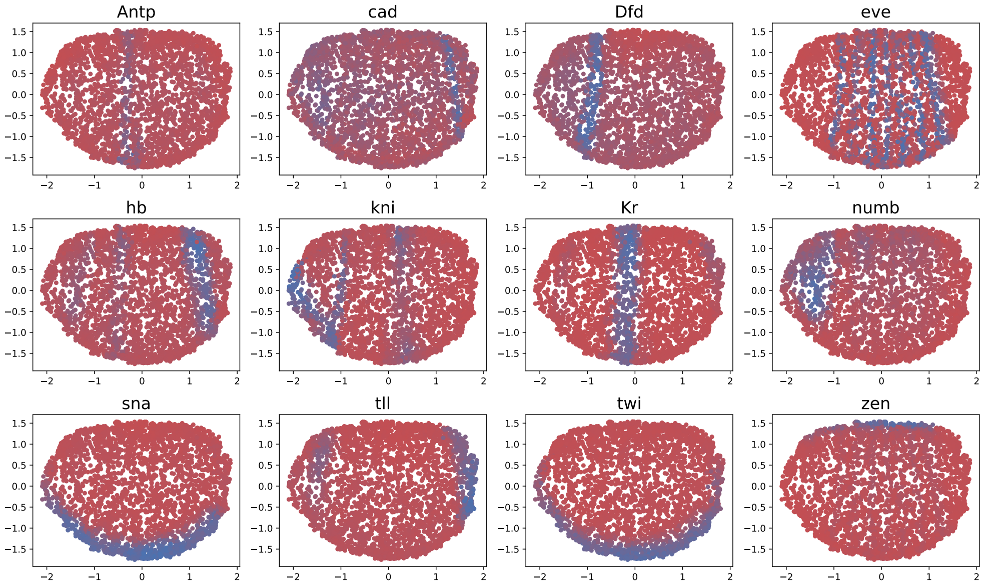

**Fig. 9 Reconstruction results of spatial expression patterns of specific genes in *Drosophila* embyro experiment by iSORT.**

435 ## 3.10 Supplementary Fig. 10: Top SOGs in human artery
436 dataset.

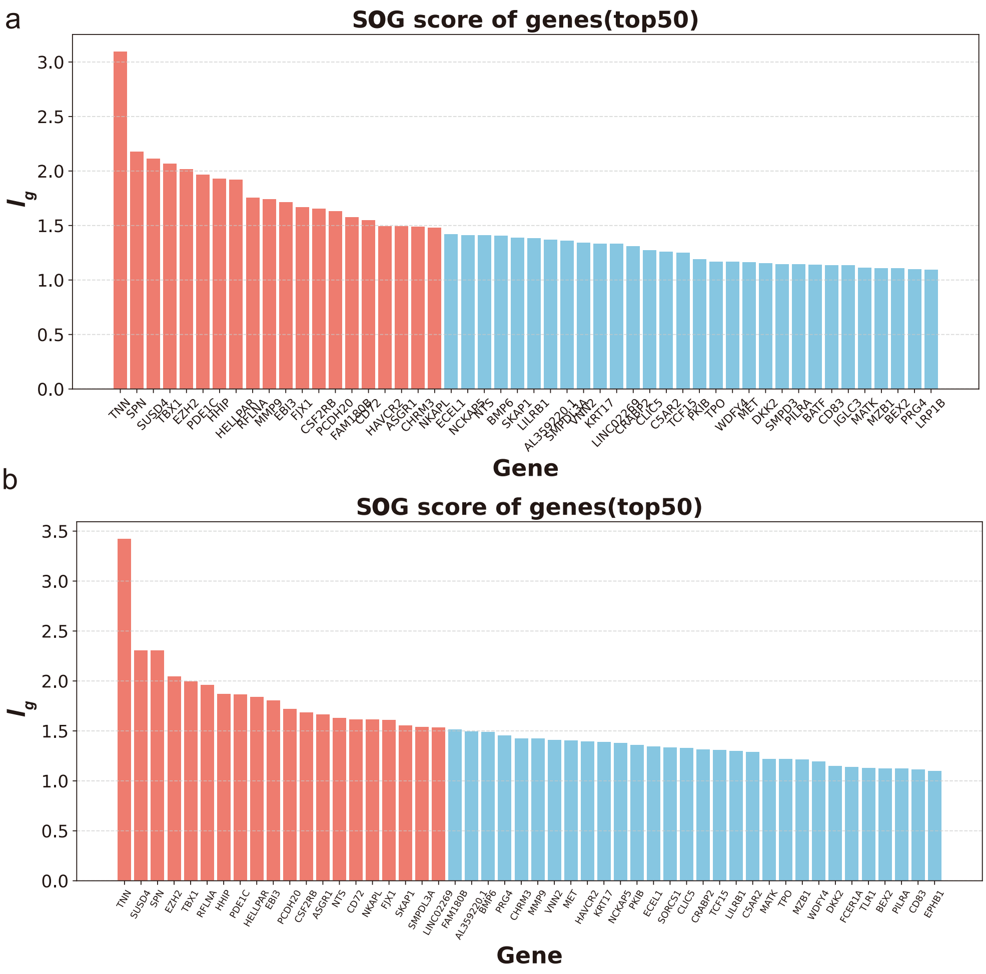

**Fig. 10 Top SOGs in human artery dataset.** (**a**) Top 50 SOGs in diseased arteries with atherosclerosis. (The 20 highest scoring genes are shown in red.) (**b**) Top 50 SOGs in normal arteries. (The 20 highest scoring genes are shown in red.)

#### 3.11 Supplementary Fig. 11: The top 20 GO terms of the 50 SOGs found by iSORT in diseased arteries with atherosclerosis.

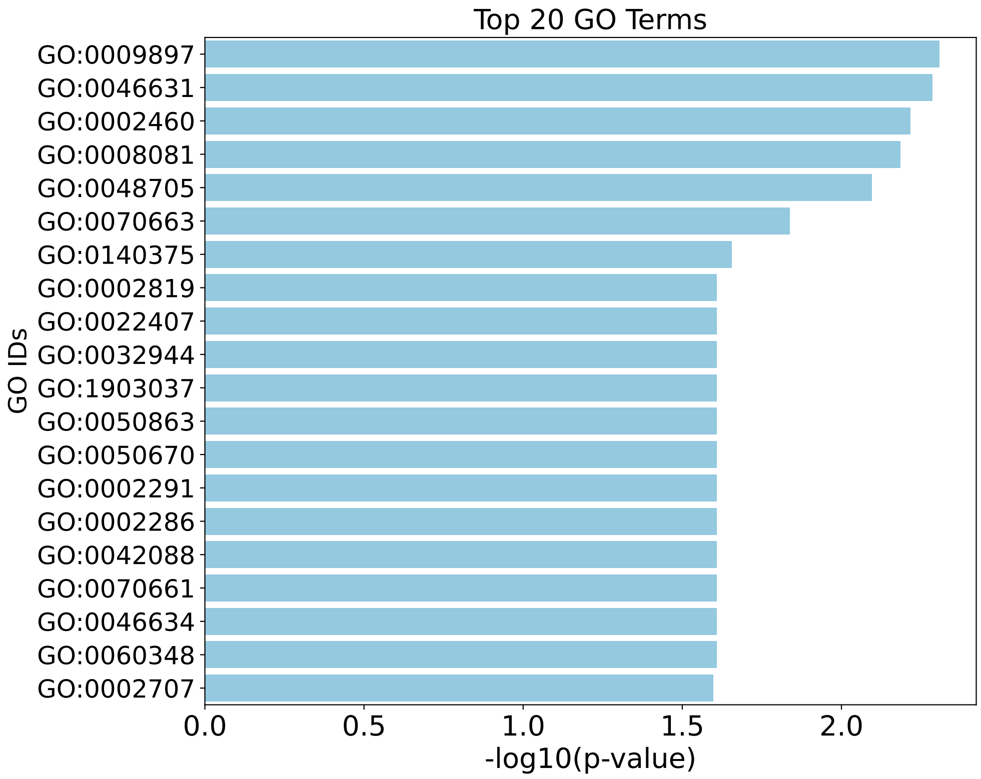

**Fig. 11** **The top 20 GO terms of the 50 SOGs found by iSORT in diseased arteries with atherosclerosis.**

#### 3.12 Supplementary Fig. 12: The top 20 GO terms of the 50 SOGs found by iSORT in normal arteries.

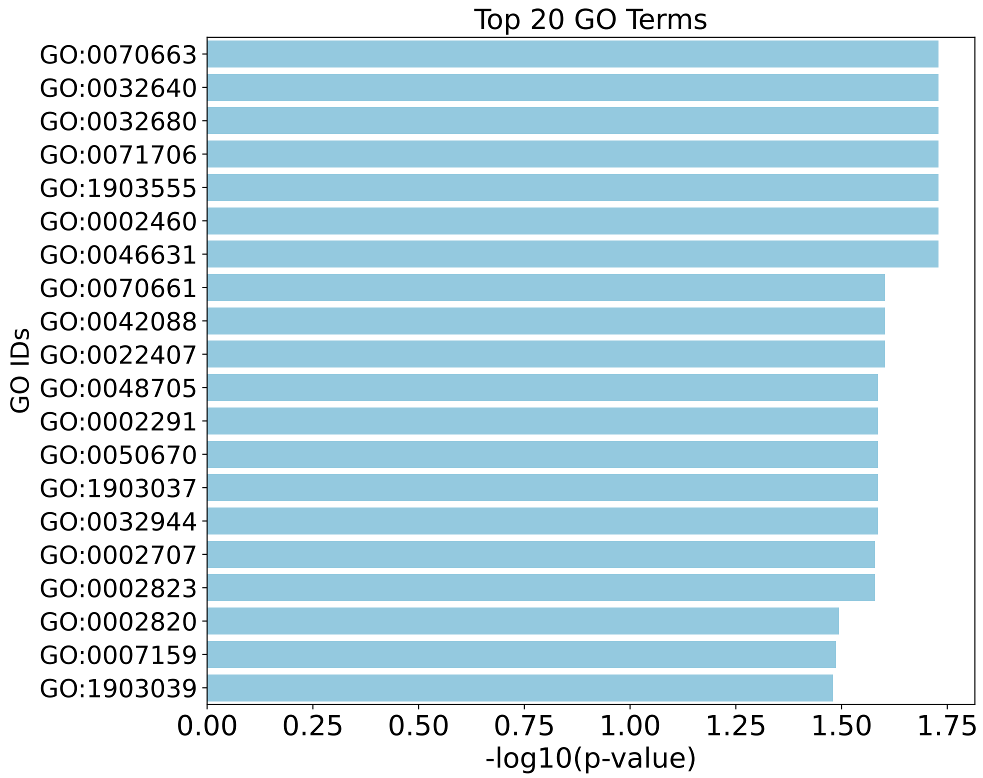

**Fig. 12** **The top 20 GO terms of the 50 SOGs found by iSORT in normal arteries.**

#### 3.13 Supplementary Fig. 13: Reconstruction results of the diseased arteries with atherosclerosis.

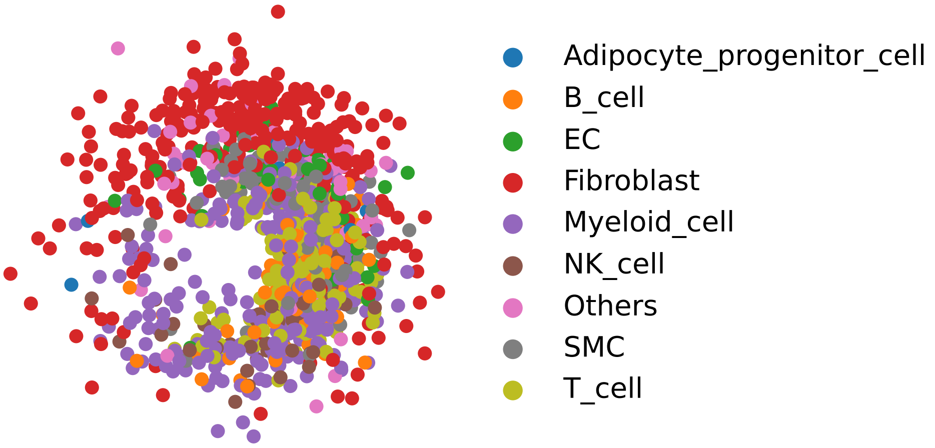

**Fig. 13 Reconstruction results of the diseased arteries with atherosclerosis.** The figure contains full cell types included in the diseased-artery dataset.

#### 3.14 Supplementary Fig. 14: Reconstruction results of normal arteries.

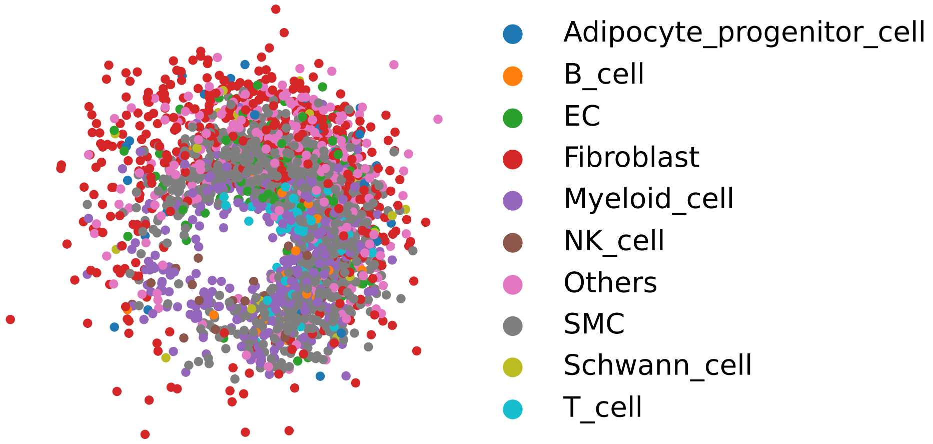

**Fig. 14 Reconstruction results of normal arteries.** The figure contains full cell types included in the normal-artery dataset.

#### 3.15 Supplementary Fig. 15: Spatial velocity of human developmental heart dataset.

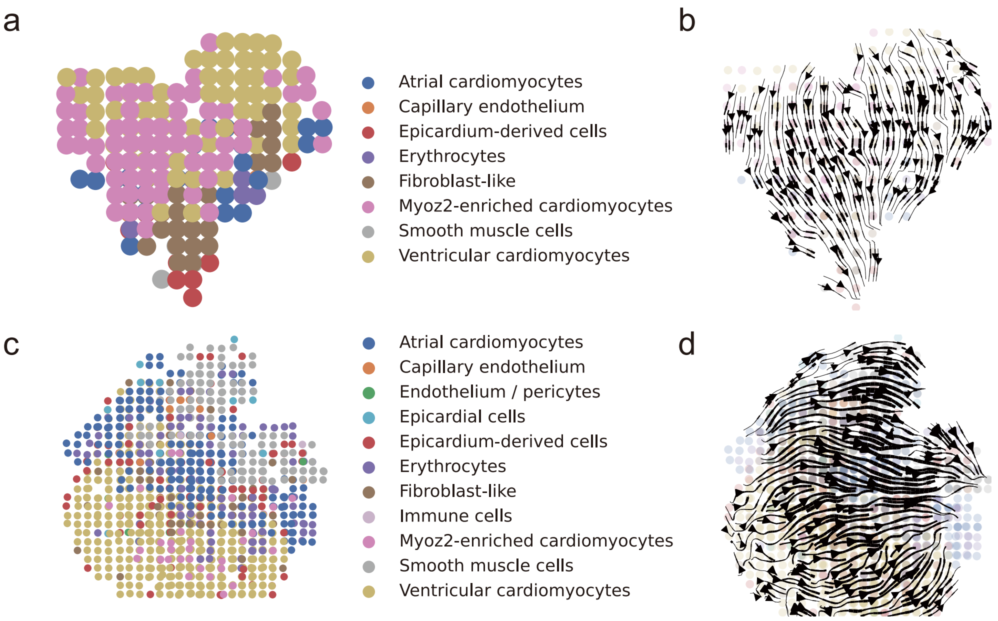

**Fig. 15 Spatial velocity of human developmental heart dataset.** (**a**) Spatial transcriptomics (ST) of a human developing heart at 4.5-5 post-conception week (PCW). (**b**) Spatial velocity on the human developing heart at 4.5-5 PCW visualized by iSORT. The results are consistent with the development of true heart. (**c**) ST of a human developing heart at 6.5 PCW. (**d**) Spatial velocity on the human developing heart at 6.5 PCW visualized by iSORT. The results are consistent with the development of true heart.

### References

[1] Sun, Q., Chattopadhyay, R., Panchanathan, S. & Ye, J. A two-stage weighting framework for multi-source domain adaptation. *Advances in neural information processing systems* **24** (2011).

[2] Qian, J. *et al.* Reconstruction of the cell pseudo-space from single-cell RNA sequencing data with scSpace. *Nature Communications* **14**, 2484 (2023).

[3] Biancalani, T. *et al.* Deep learning and alignment of spatially resolved single-cell transcriptomes with Tangram. *Nature Methods* **18**, 1352–1362 (2021).

[4] Nitzan, M., Karaiskos, N., Friedman, N. & Rajewsky, N. Gene expression cartography. *Nature* **576**, 132–137 (2019).

[5] Zhang, Q. *et al.* Leveraging spatial transcriptomics data to recover cell locations in single-cell RNA-seq with CeLEry. *Nature Communications* **14**, 4050 (2023).

[6] Wei, R. *et al.* Spatial charting of single-cell transcriptomes in tissues. *Nature Biotechnology* **40**, 1190–1199 (2022).

[7] Svensson, V., Teichmann, S. A. & Stegle, O. Spatialde: identification of spatially variable genes. *Nature Methods* **15**, 343–346 (2018).

[8] Getis, A. & Ord, J. K. The analysis of spatial association by use of distance statistics. *Geographical Analysis* **24**, 189–206 (1992).

[9] Palla, G. *et al.* Squidpy: a scalable framework for spatial omics analysis. *Nature Methods* **19**, 171–178 (2022).

[10] Maynard, K. R. *et al.* Transcriptome-scale spatial gene expression in the human dorsolateral prefrontal cortex. *Nature Neuroscience* **24**, 425–436 (2021).

[11] Lohoff, T. *et al.* Integration of spatial and single-cell transcriptomic data elucidates mouse organogenesis. *Nature Biotechnology* **40**, 74–85 (2022).

[12] Tasic, B. *et al.* Adult mouse cortical cell taxonomy revealed by single cell transcriptomics. *Nature Neuroscience* **19**, 335–346 (2016).

[13] Berkeley Drosophila Transcription Network Project. URL http://bdtnp.lbl.gov/. Available at: http://bdtnp.lbl.gov/.

[14] Luengo Hendriks, C. L. *et al.* Three-dimensional morphology and gene expression in the Drosophilablastoderm at cellular resolution I: data acquisition pipeline. *Genome Biology* **7**, 1–21 (2006).

[15] Asp, M. *et al.* A spatiotemporal organ-wide gene expression and cell atlas of the developing human heart. *Cell* **179**, 1647–1660 (2019).

[16] Scherberich, A. *et al.* Tenascin-W is found in malignant mammary tumors, promotes alpha8 integrin-dependent motility and requires p38MAPK activity for BMP-2 and TNF-alpha induced expression in vitro. *Oncogene* **24**, 1525–1532 (2005).

[17] Tucker, R. P., Ferralli, J., Schittny, J. C. & Chiquet-Ehrismann, R. Tenascin-C and tenascin-W in whisker follicle stem cell niches: possible roles in regulating stem cell proliferation and migration. *Journal of Cell Science* **126**, 5111–5115 (2013).

[18] Neidhardt, J., Fehr, S., Kutsche, M., Löhler, J. & Schachner, M. Tenascin-N: characterization of a novel member of the tenascin family that mediates neurite repulsion from hippocampal explants. *Molecular and Cellular Neuroscience* **23**, 193–209 (2003).

[19] Meloty-Kapella, C. V., Degen, M., Chiquet-Ehrismann, R. & Tucker, R. P. Effects of tenascin-W on osteoblasts in vitro. *Cell and Tissue Research* **334**, 445–455 (2008).

[20] Zhou, A.-X., Hartwig, J. H. & Akyürek, L. M. Filamins in cell signaling, transcription and organ development. *Trends in Cell Biology* **20**, 113–123 (2010).

[21] Nakamura, F., Stossel, T. P. & Hartwig, J. H. The filamins: organizers of cell structure and function. *Cell Adhesion & Migration* **5**, 160–169 (2011).

[22] Campbell, I. D. Studies of focal adhesion assembly. *Biochemical Society Transactions* **36**, 263–266 (2008).

[23] Bandaru, S. *et al.* Targeting filamin A reduces macrophage activity and atherosclerosis. *Circulation* **140**, 67–79 (2019).

[24] Seo, W. & Ziltener, H. J. CD43 processing and nuclear translocation of CD43 cytoplasmic tail are required for cell homeostasis. *Blood, The Journal of the American Society of Hematology* **114**, 3567–3577 (2009).

[25] Mody, P. D. *et al.* Signaling through CD43 regulates CD4 T-cell trafficking. *Blood, The Journal of the American Society of Hematology* **110**, 2974–2982 (2007).

[26] Serrador, J. M. *et al.* CD43 interacts with moesin and ezrin and regulates its redistribution to the uropods of T lymphocytes at the cell-cell contacts. *Blood, The Journal of the American Society of Hematology* **91**, 4632–4644 (1998).

[27] McEvoy, L. M., Jutila, M. A., Tsao, P. S., Cooke, J. P. & Butcher, E. C. Anti-CD43 inhibits monocyte-endothelial adhesion in inflammation and atherogenesis. *Blood, The Journal of the American Society of Hematology* **90**, 3587–3594 (1997).

[28] Chittock, E. C., Latwiel, S., Miller, T. C. & Müller, C. W. Molecular architecture of polycomb repressive complexes. *Biochemical Society Transactions* **45**, 193–205 (2017).

[29] Schmitges, F. W. *et al.* Histone methylation by PRC2 is inhibited by active chromatin marks. *Molecular Cell* **42**, 330–341 (2011).

[30] Schuettengruber, B. & Cavalli, G. Recruitment of polycomb group complexes and their role in the dynamic regulation of cell fate choice. *Development* **136**, 3531–3542 (2009).

[31] Greißel, A. *et al.* Histone acetylation and methylation significantly change with severity of atherosclerosis in human carotid plaques. *Cardiovascular Pathology* **25**, 79–86 (2016).

[32] Meng, X.-D. *et al.* Knockdown of GAS5 inhibits atherosclerosis progression via reducing EZH2-mediated ABCA1 transcription in ApoE-/- mice. *Molecular Therapy-Nucleic Acids* **19**, 84–96 (2020).
